# The *Ralstonia solanacearum* E3 ligase effector RipV1 targets subfamily IXb receptor-like cytoplasmic kinases that negatively regulate immunity in *Nicotiana benthamiana*

**DOI:** 10.1101/2025.07.21.665888

**Authors:** Jihyun Choi, Thakshila Dharmasena, Yoonyoung Lee, Jacqueline Monaghan, Cécile Segonzac

**Author notes:** For correspondence; https://orcid.org/0000-0002-5537-7556. Present address: Institute of Plant and Microbial Biology, Zurich-Basel Plant Science Center, University of Zurich, 8008 Zurich, Switzerland (Yoonyoung Lee).

## Abstract

Plants detect microbe-associated molecular patterns from pathogens via plasma membrane-localized receptors which activate multiple signaling cascades that lead to pattern-triggered immunity (PTI). Receptor-like cytoplasmic kinases (RLCKs) are essential hubs of plant immune signaling, associating with receptors and intracellular proteins through phosphorylation events. As a consequence, RLCKs have emerged as common targets of pathogen effectors. To improve our knowledge on Solanaceae responses to the bacterial wilt pathogen *Ralstonia solanacearum*, we conducted a yeast two-hybrid screen between tomato RLCKs and effectors conserved in *R. solanacearum* Korean isolates. Several members of RLCK subfamily IXb, which contain a ubiquitin-ligase plant U-box domain in addition to the kinase domain, interacted with RipV1, an effector containing a novel E3 ligase domain (NEL). *In vitro* assays revealed that RLCK-IXb-1 displayed ubiquitin-ligase activity but no detectable kinase activity. RipV1 could trans- ubiquitinate RLCK-IXb-1 *in vitro* and promote its stability *in planta*. Using virus-induced gene silencing of RLCK-IXb homologs in *Nicotiana benthamiana*, we could further show that several RLCK-IXb proteins act as negative regulators of early PTI signaling. RipV1 was previously reported to contribute to *R. solanacearum* virulence in potato and to elicit cell death in an E3 ligase activity-dependent manner in *N. benthamiana*. Here we show that RipV1-induced cell death occurred in plants impaired for effector recognition but could be suppressed by over-expression of RLCK-IXb-1, suggesting that this response is related to the virulence function of RipV1. Altogether, our work identifies possible substrates of an NEL effector and underlines the complex roles of RLCKs in plant immune signaling.

## INTRODUCTION

*Ralstonia solanacearum* is a globally destructive soil-borne pathogen infecting Solanaceous crops (Ahmed et al., 2022). *R. solanacearum* delivers over 70 type III secreted effectors, Ralstonia injected proteins (Rips), into host cells to suppress plant immunity and promote virulence (Landry et al., 2020; Peeters et al., 2013). A subset of *R. solanacearum* effectors, including GAxxL motif-containing (GALA)/RipG and Novel E3 ligase (NEL) effectors, are involved in ubiquitination processes in host plant cells and contribute to virulence (Lei et al., 2020; Pedro- Jové et al., 2021; Remigi et al., 2011). GALA/RipG effectors contain F-box motifs that facilitate their interaction with Skp-Cullin-F-box (SCF) ubiquitin-ligase complexes in *Arabidopsis thaliana* (Angot et al., 2006). In contrast, NEL effectors carry a distinct ubiquitin ligase domain that functions independently of the canonical SCF machinery and are divergent from previously characterized bacterial E3 ligases (Bullones-Bolaños et al., 2022; Rohde et al., 2007). Four *R. solanacearum* NEL domain-containing effectors, RipAR, RipAW, RipV1 and RipV2, have been reported to auto-ubiquitinate and alter host immune responses (Cheng et al., 2021; Cheng et al., 2024; Nakano et al., 2017; Ouyang et al., 2023; Qi et al., 2024). When heterologously expressed in *Nicotiana benthamiana*, the four NEL effectors suppress plant defense signaling triggered by the microbe-associated molecular pattern (MAMP) bacterial flagellin epitope flg22 in an E3 ligase-dependent manner (Cheng et al., 2021; Cheng et al., 2024; Nakano et al., 2017). Plant intracellular nucleotide-binding leucine-rich repeat receptors (NLRs) detect the presence or activity of pathogen effectors and activate a conserved signaling network leading to effector-triggered immunity (ETI) and often a form of programed cell death termed the hypersensitive response (Jones et al., 2024). In *N. benthamiana*, NEL effectors RipAW, RipV1 and RipV2 also elicit weak to robust cell death in an E3 ligase-dependent manner, suggesting that NEL effector activity could dramatically impair host cell functions or alternatively, could be monitored by the plant immune system (Cheng et al., 2021; Cheng et al., 2024; Niu et al., 2021).

Most NLRs are regulated by a ubiquitin-ligase complex containing SUPPRESSOR OF THE G2 ALLELE OF SKP1 (SGT1); *SGT1* silencing suppresses NLR-dependent cell death and can also impair protein accumulation in the context of Agrobacterium-mediated expression (Shirasu, 2009; Yu et al., 2019). The Toll-interleukin 1-receptor (TIR) domain containing NLRs (TNLs) require the helper NLR N REQUIREMENT GENE 1 (NRG1) and lipase-like proteins ENHANCED DISEASE SUSCEPTIBILITY 1 (EDS1), PHYTOALEXIN DEFICIENT 4 (PAD4), and SENESCENCE ASSOCIATED GENE 101 (SAG101) to signal ETI (Dongus and Parker, 2021). A recent study revealed that the *R. solanacearum* NEL effector RipV2 destabilizes NRG1 and the EDS1/SAG101 complex through ubiquitination and 26S proteasome degradation, hence inhibiting bacterial wilt resistance in tomato (Qi et al., 2024). NEL effectors could also affect host basal defense signaling, as RipAW can ubiquitinate and destabilize the pattern-recognition receptor complex formed by the leucine-rich repeat receptor-like kinase (LRR-RLK) FLAGELLIN SENSING 2 (FLS2) and the receptor-like cytoplasmic kinase (RLCK) BOTRYTIS INDUCED KINASE 1 (BIK1) in Arabidopsis (Sun et al., 2024). Identifying other substrates of NEL effectors is therefore necessary to understand their contribution to virulence.

Protein phosphorylation acts as a molecular switch in signaling pathways. This post-translational modification is mediated by protein kinases such as RLKs and RLCKs, mitogen-activated protein kinases, and calcium-dependent protein kinases (Bredow and Monaghan, 2019; Jagodzik et al., 2018; Liu et al., 2024; Sun and Zhang, 2022; Valmonte et al., 2014). RLCKs are categorized in subfamilies based on the conservation of the kinase domain and play a central role in transducing signals from cell surface-localized RLKs to intracellular signaling components (Hailemariam et al., 2024; He et al., 2018). In the plant immune system, the subfamily VII RLCK BIK1 functions downstream of MAMP-recognition receptors and positively regulates early immune responses such as the production of reactive oxygen species (ROS) through the phosphorylation of the respiratory burst oxidase RbohD (Li et al., 2014; Lu et al., 2010; Kadota et al., 2014). RLCKs act redundantly as drivers of immune signaling pathways and are targeted by diverse bacterial effectors (Sun and Zhang, 2020). For example, the *Xanthomonas campestris* pv. *campestris* effector AvrAC uridylylates and inhibits BIK1 kinase activity, leading to suppression of plant defense responses (Feng et al., 2012). In addition, *Pseudomonas syringae* effectors HopZ1a, AvrB, and AvrPphB can also modify multiple members of the RLCK-VII subfamily to impair plant defense (Bastedo et al., 2019; Liu et al., 2011; Russell et al., 2015; Shao et al., 2003; Zhang et al., 2010). RLCKs could thus represent potential targets for *R. solanacearum* effectors. Similar to phosphorylation, ubiquitination plays a key role in plant immune signaling (Trujillo and Shirasu, 2010; Langin et al., 2023). The ubiquitination process is mediated by a cascade of enzymatic activities involving first a ubiquitin-activating enzyme (E1), then a ubiquitin-conjugating enzyme (E2), and finally a ubiquitin-ligase (E3). While E2 enzymes determine the type of ubiquitin linkage (Trujillo, 2017), E3 ligases confer substrate specificity (Toma-Fukai and Shimizu, 2021). The fate of ubiquitinated proteins depend on the type of linkage, with polyubiquitination generally leading to endocytosis or degradation by the 26S proteasome, while monoubiquitination can enhance stability or change subcellular localization (Zhou and Zeng, 2017). In plants, a family of E3 ligases harbor a U-box domain and are collectively referred to as plant U-box proteins (PUBs). PUBs are divided into 10 classes based on the presence of additional domains (Azevedo et al., 2001; Trenner et al., 2022; Wiborg et al., 2008). PUBs often function as homeostatic regulators of plant immunity (Tenner et al., 2022). In Arabidopsis, BIK1 homeostasis is regulated by PUB25 and PUB26 through ubiquitination (Dias et al., 2022; Fu et al., 2024; Wang et al., 2018). PUBs are also targeted by pathogen effectors. In rice, the *X. oryzae* pv. *oryzae* effector XopP interacts with and suppresses the E3 ligase activity of OsPUB44, promoting pathogen infection (Ishikawa et al., 2014). Similarly, two *R. solanacearum* effectors can modify host PUB activity. RipAC targets PUB4, enhancing BIK1 degradation and suppressing MAMP-triggered immunity (Yu et al., 2022). RipAV also interacts with PUBs that ubiquitinate BIK1, promoting BIK1 degradation to suppress immune signaling (Rufian et al., 2025).

Here, we identified a subset of subfamily IXb RLCKs from tomato that contains an integrated PUB domain and interacts with *R. solanacearum* NEL effector RipV1. We could show that *in vitro* RLCK-IXb-1 is an active E3 ligase but does not display detectable kinase activity. While we could not ascertain whether RLCK-IXb-1 ubiquitinates RipV1, we did observe that RipV1 ubiquitinates RLCK-IXb-1 *in vitro* and promotes its stability *in planta*. We further demonstrated that RLCK-IXb proteins can act as negative regulators of early immune responses and RipV1-triggered cell death in *N. benthamiana*. Overall, our work identifies possible substrates of a NEL effector and underlines the complex roles of RLCKs in plant immune signaling.

## RESULTS

### RipV1 interacts with several RLCK-IXb proteins via its N-terminal domain

RLCKs contribute to phospho-relay in plant immune signaling and are targets of bacterial effectors (Lu et al., 2010; Wang et al., 2015; Zhang et al., 2010). We hypothesized that *R. solanacearum* effectors could target RLCKs. Using a LexA-based yeast two-hybrid system (Y2H), we screened 40 effectors conserved in *R. solanacearum* Korean isolates for interaction with 127 tomato RLCKs (Prokchorchik et al., 2020; Sakamoto et al., 2012). Among positive interactions, we found that the NEL-effector RipV1 could interact with 4 members of the RLCK-IXb subfamily (RLCK-IXb-1, -2, -3, and -9; Figure 1A). The RLCK-IXb subfamily is notable because of the presence of an integrated PUB domain C-terminal to the kinase domain (Sakamoto et al., 2012; Trenner et al., 2022; Figures S1 and S2). To refine the interaction interface using Y2H, we designed fragments corresponding to domains in RipV1 and RLCK-IXb-1 as predicted from structural models generated by AlphaFold 3. RipV1 protein consists of an N-terminal domain structured by α-helices (1-374 aa) and the C-terminal NEL domain (375-685 aa) (Figures 1B, S1A and S1B). Only the N-terminal domain of RipV1 could interact with full length RLCK-IXb-1 (Figure 1B), indicating that the NEL domain was dispensable for the interaction. The RLCK-IXb-1 protein includes an N-terminal universal stress protein domain (USP; 1-208 aa), a structural maintenance of chromatin domain (Smc; 360-520 aa) that corresponds to a long α-helix in the 3D model, the protein kinase domain (Kinase; 521-814 aa) and the C-terminal PUB domain (PUB; 815-894 aa) (Figures 1C, S1C and S1D). Only the fragments containing the Smc domain could interact with the RipV1 N-terminal domain (Figure 1C). Together, these results suggest that the N-terminal domain of RipV1 and the Smc domain of RLCK-IXb-1 mediate the interaction between the two proteins.

**Figure 1.**
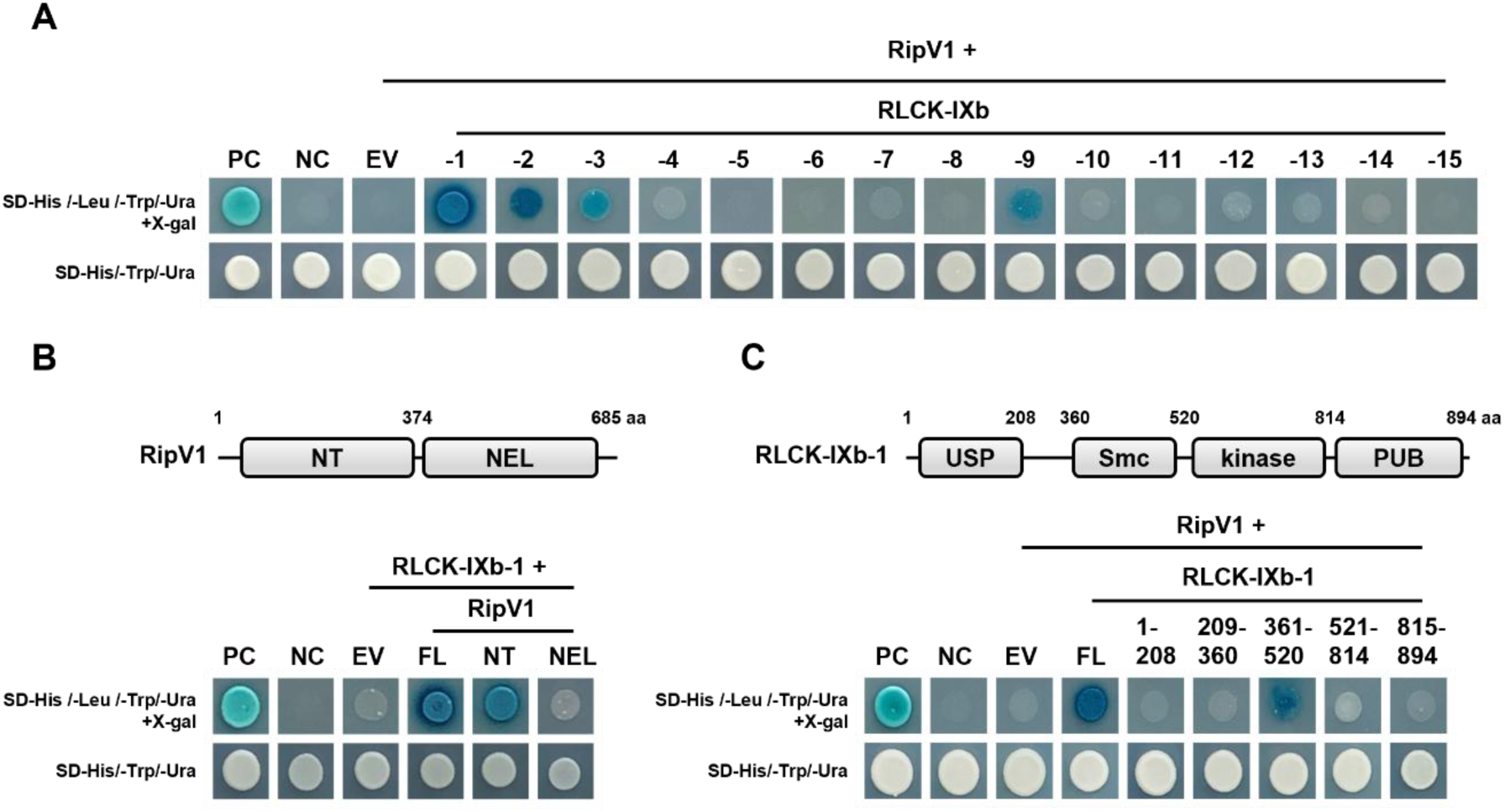
RipV1 interacts with tomato RLCK-IXb via its N-terminal domain. (**A**) RipV1 interacted with several members of the tomato RLCK subfamily IXb in yeast two-hybrid. Yeast strain EGY48 carrying RipV1-FLAG fused to the LexA DNA-binding domain in the pLexA vector was mated with a set of RFY206 strains, each carrying a different tomato RLCK-IXb gene fused to the B42 activation domain in the pB42-AD vector. Mated cells were spotted on selective media [SD- His/-Leu /-Trp/-Ura containing 5-Bromo-4-Chloro-3-Indolyl β-D-galactopyranoside (X-gal)] and photographed. EDS1–PAD4 interaction was used as a positive control (PC). Mating of cells carrying the pLexA and pB42-AD empty vectors was used as a negative control (NC). (**B**) The N- terminal domain of RipV1 was required for the interaction with RLCK-IXb-1. RipV1 was divided into two domains based on structural analysis (FL, full-length, 1-685 aa; NT, N-terminal, 1-374 aa; NEL, NEL domain, 375-685 aa) and tested for interaction with RLCK-IXb-1. (**C**) The Smc domain of RLCK-IXb-1 interacted with the N-terminal domain of RipV1. RLCK-IXb-1 was divided into five domains based on structure analysis (FL, full length, 1-894 aa; USP, Universal Stress Protein, 1-208 aa; flexible region, 209-360 aa; Smc, structural maintenance of chromatin, 361- 520 aa; Kinase, 521-814 aa; PUB, 815-894 aa) and tested for interaction with the RipV1 N- terminal domain. Predicted structures for RipV1 and RLCK-IXb-1 are shown in Figure S1.

To confirm the interaction *in planta*, we conducted a co-immunoprecipitation assay in *N. benthamiana* tissues expressing RipV1 fused to the yellow fluorescent protein (YFP) and RLCK-IXb-1 fused to 6 hemagglutinin tags (HA). RLCK-IXb-1-HA co-immunoprecipitated with RipV1-YFP but not with the green fluorescent protein (GFP) control (Figure 2A). We further characterized the subcellular localization of RipV1 and RLCK-IXb-1 in plant cells. Confocal imaging revealed that RipV1-YFP localizes to the cell periphery in agreement with a previous report (Cheng et al., 2024) and co-localizes with the plasma membrane marker FLS2-mCherry (Figures 2B and S3A). The RLCK-IXb-1-YFP signal was similarly observed at the cell periphery (Figure S3B). Some RLCKs have putative N-terminal myristoylation and/or palmitoylation sites that facilitate localization to the plasma membrane (Veronese et al., 2006). Although no myristoylation site was predicted in RLCK-IXb-1, several predicted palmitoylation sites may contribute to the peripheral localization (Table S1). RLCK-IXb-1 also showed some accumulation in the nucleus, which could not be attributed to cleaved YFP on immunoblot (Figure S3B and S3C). In cells co-expressing RipV1-YFP and RLCK-IXb-1 fused to the cyan fluorescent protein (CFP), a clear overlap of the fluorescence signals could be observed at the cell periphery (Figure 2B). These results validate the interaction between RipV1 and RLCK-IXb-1 *in planta* and suggest that their association occurs in the proximity of the plasma membrane.

**Figure 2.**
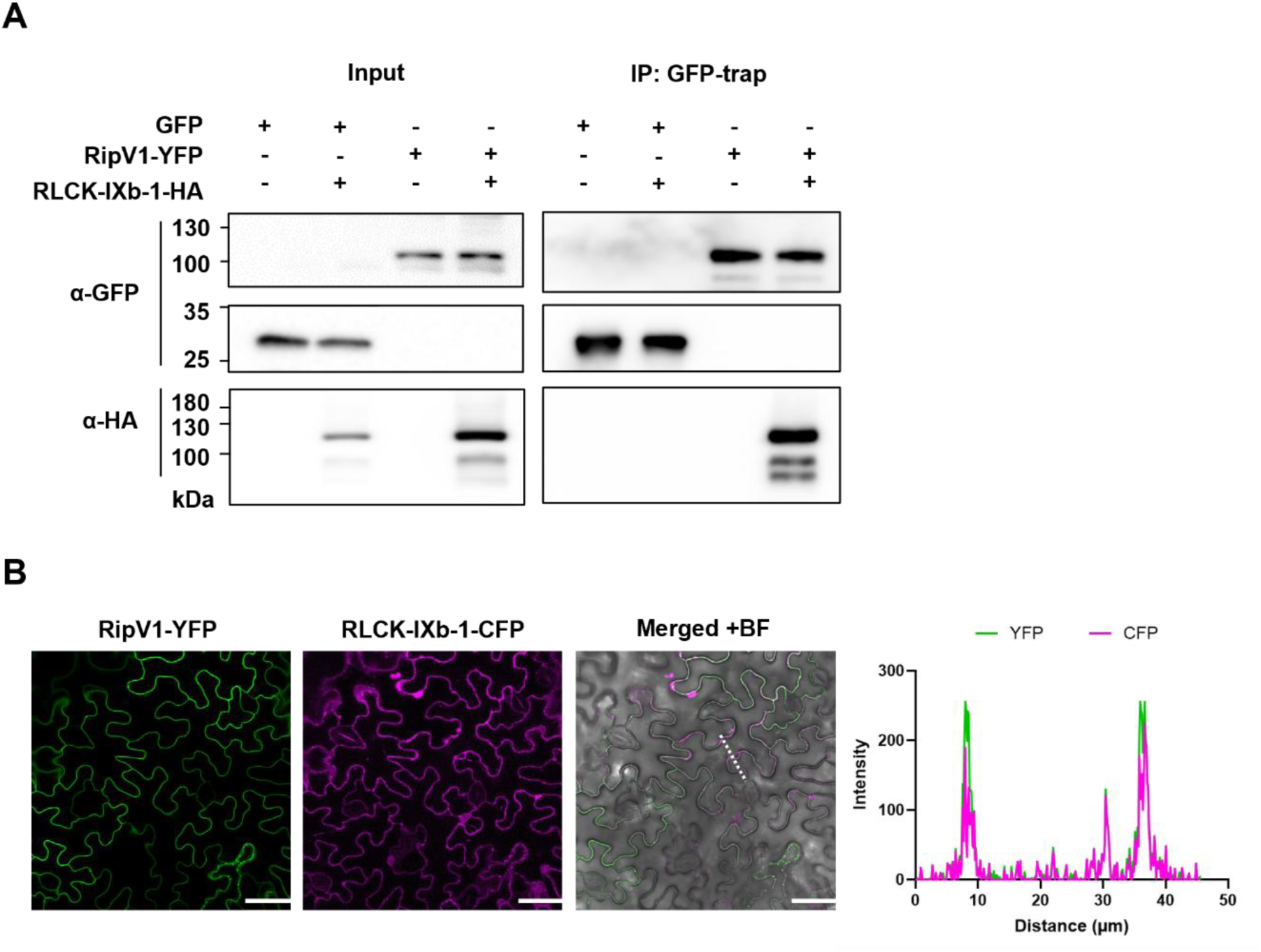
RipV1 associates with RLCK-IXb-1 *in planta*. (**A**) Green fluorescent protein (GFP) or RipV1 C-terminally fused to yellow fluorescent protein (YFP) were co-expressed with RLCK-IXb- 1 C-terminally tagged with hemagglutinin (HA) in *N. benthamiana*. Leaf tissues were harvested 45 hours post infiltration, and total proteins were extracted. Total protein extracts were incubated with GFP-trap beads for immunoprecipitation. Total extract proteins (input) and immunoprecipitated (IP) proteins were probed with anti-GFP or anti-HA antibodies. (**B**) RipV1 and RLCK-IXb-1 co-localized at the cell periphery. RipV1-YFP and RLCK-IXb-1 fused to C-terminal cyan fluorescent protein (CFP) were co-expressed in *N. benthamiana*. Subcellular localization of RipV1-YFP and RLCK-IXb-1-CFP was observed by confocal microscopy. Fluorescence images were merged with the brightfield (BF) image. Scale bars represent 50 µm. Fluorescence intensity of YFP and CFP across the section indicated by the white dotted line is shown on the right.

### RLCK-IXb-1 is a functional E3 ligase but lacks detectable kinase activity *in vitro*

RLCK-IXb-1 belongs to PUB class VI containing both protein kinase and PUB domains (Figures S1C, S1D and S2) (Sakamoto et al., 2012; Trenner et al., 2022). To determine RLCK-IXb-1 enzymatic activities, recombinant RLCK-IXb-1 and predicted catalytic dead mutants in the PUB domain (RLCK-IXb-1^C830A^) and the kinase domain (RLCK-IXb-1^D659A^), all tagged with 3xFLAG were purified. For *in vitro* ubiquitination assays, recombinant proteins were incubated with ubiquitin-HA, E1 activating (UBE1) and E2 conjugating (UbcH5B) enzymes. High molecular weight polyubiquitination smears were detected on RLCK-IXb-1 (Figure 3A), indicative of auto- ubiquitination. In contrast, no auto-ubiquitination could be detected on the PUB mutant RLCK-IXb-1^C830A^, in which a conserved cysteine in the PUB domain is mutated to alanine (Lu et al. 2011). Together, these results indicate that RLCK-IXb-1 possesses catalytic E3 ligase activity *in vitro*.

**Figure 3.**
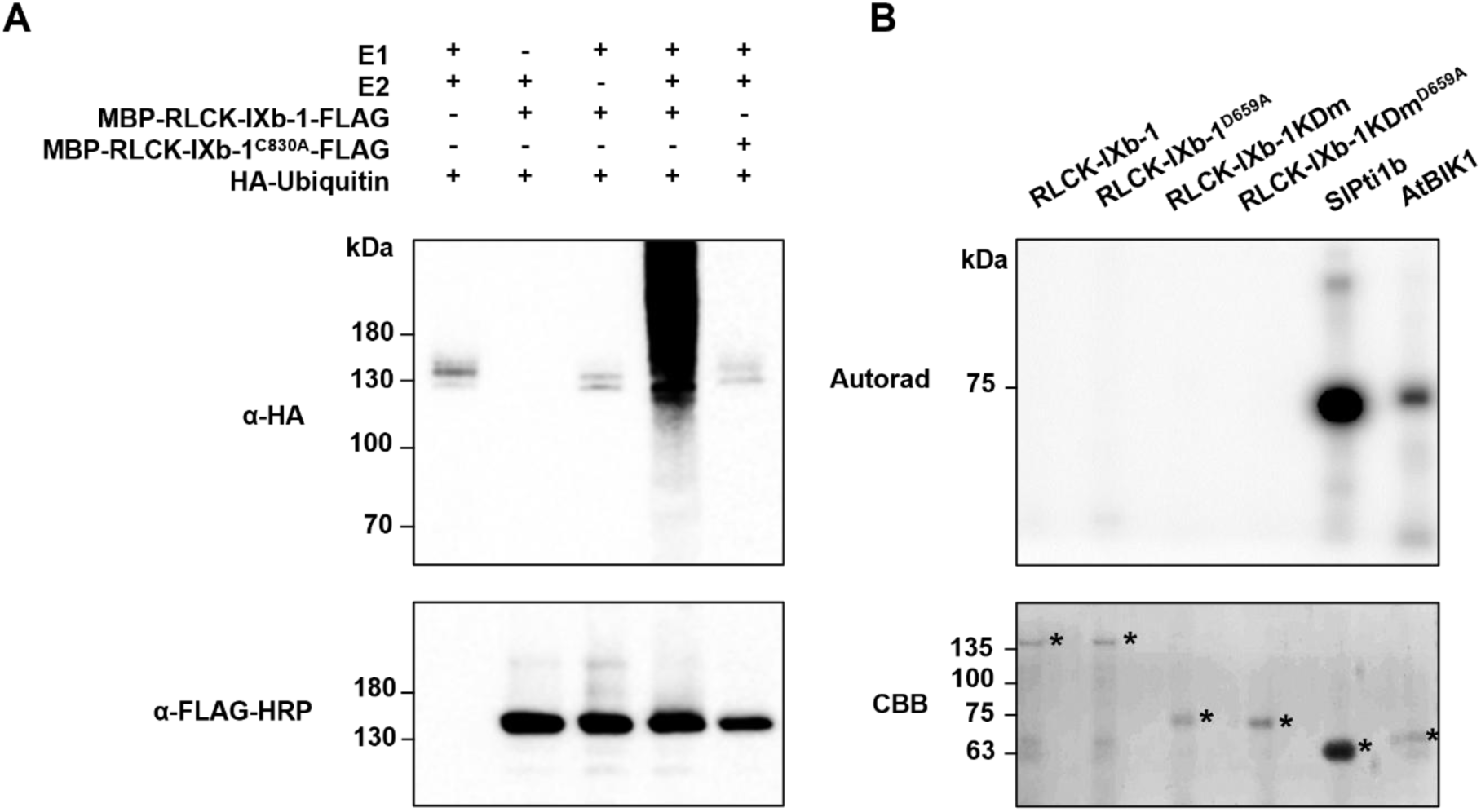
RLCK-IXb-1 is a functional E3 ligase but lacks detectable kinase activity *in vitro*. (**A**) Auto-ubiquitination of RLCK-IXb-1 *in vitro*. FLAG-tagged RLCK-IXb-1 and E3 ligase-dead mutant (RLCK-IXb-1^C830A^) recombinant proteins were incubated with HA-Ubiquitin in the presence or absence of human E1 (UBE1) and E2 (UbcH5B). Reaction mixtures were separated by SDS-PAGE and detected by immunoblot assay using anti-HA or anti-FLAG-HRP antibodies. (**B**) *In vitro* kinase assay of RLCK-IXb-1 derivatives. RLCK-IXb-1, kinase-dead mutant (RLCK-IXb-1^D659A^), isolated kinase domain (RLCK-IXb-1KDm), and kinase-dead mutant of the isolated kinase domain (RLCK- IXb-1KDm^D659A^) were incubated with kinase reaction buffer containing γ^32^P-ATP. RLCK-IXb-1KDm comprised residues 521-814, identical to the kinase domain truncation employed in yeast two- hybrid. Phosphorylation was detected by autoradiography (Autorad) following SDS-PAGE. Coomassie Brilliant Blue (CBB) staining was performed to verify protein loading. SlPt1b and AtBIK1 were used as auto-phosphorylation controls. Asterisks indicate the positions of the recombinant proteins.

RLCK-IXb-1 harbors conserved motifs required for kinase activity, including the HRD motif essential for phosphor-transfer (Figure S4) (Boudeau et al., 2006; Eyers and Murphy, 2013). However, unlike canonical RD kinases, RLCK-IXb-1 contains a glycine (G) in place of the conserved arginine (R) within the HRD motif, classifying it as a non-RD kinase (Roux et al., 2014). Using radio-labeled ATP, we monitored the kinase activity of the recombinant full length RLCK-IXb-1 and RLCK-IXb-1^D659A^, as well as the isolated kinase domain (KDm) RLCK-IXb-1-KDm and RLCK-IXb-1KDm^D659A^ to account for possible allosteric hindrance (Figure 3B). RLCKs SlPti1b and AtBIK1 were used as positive controls for auto-phosphorylation (Gonçalves Dias et al., 2025; Lin et al., 2014; Xu et al., 2013). Auto-phosphorylation was not detected for RLCK-IXb-1 nor RLCK-IXb-1KDm, indicating that in our experimental conditions, RLCK-IXb-1 lacked a detectable kinase activity. Additionally, we co-incubated RLCK-IXb-1 with RipV1 and could not detect RipV1 trans- phosphorylation (Figure S5). Overall, *in vitro* assays indicated that RLCK-IXb-1 possesses E3 ligase activity but did not display detectable kinase activity on itself nor on RipV1.

### RipV1 ubiquitinates RLCK-IXb-1 *in vitro* and promotes RLCK-IXb-1 stability *in planta*

To test whether RLCK-IXb-1 can be a target of RipV1, we performed ubiquitination assays with recombinant RipV1 tagged with 4xMyc and RLCK-IXb-1^C830A^ tagged with 3xFLAG (Figure 4A). We first confirmed that we could detect RipV1 E3 ligase activity in our experimental conditions (Figures 4A and S6). As previously shown, RipV1 auto-ubiquitination activity was detected as a high molecular weight smear and was dependent on the conserved cysteine in the NEL domain (mutated in RipV1^C452A^) (Cheng et al., 2024).

**Figure 4.**
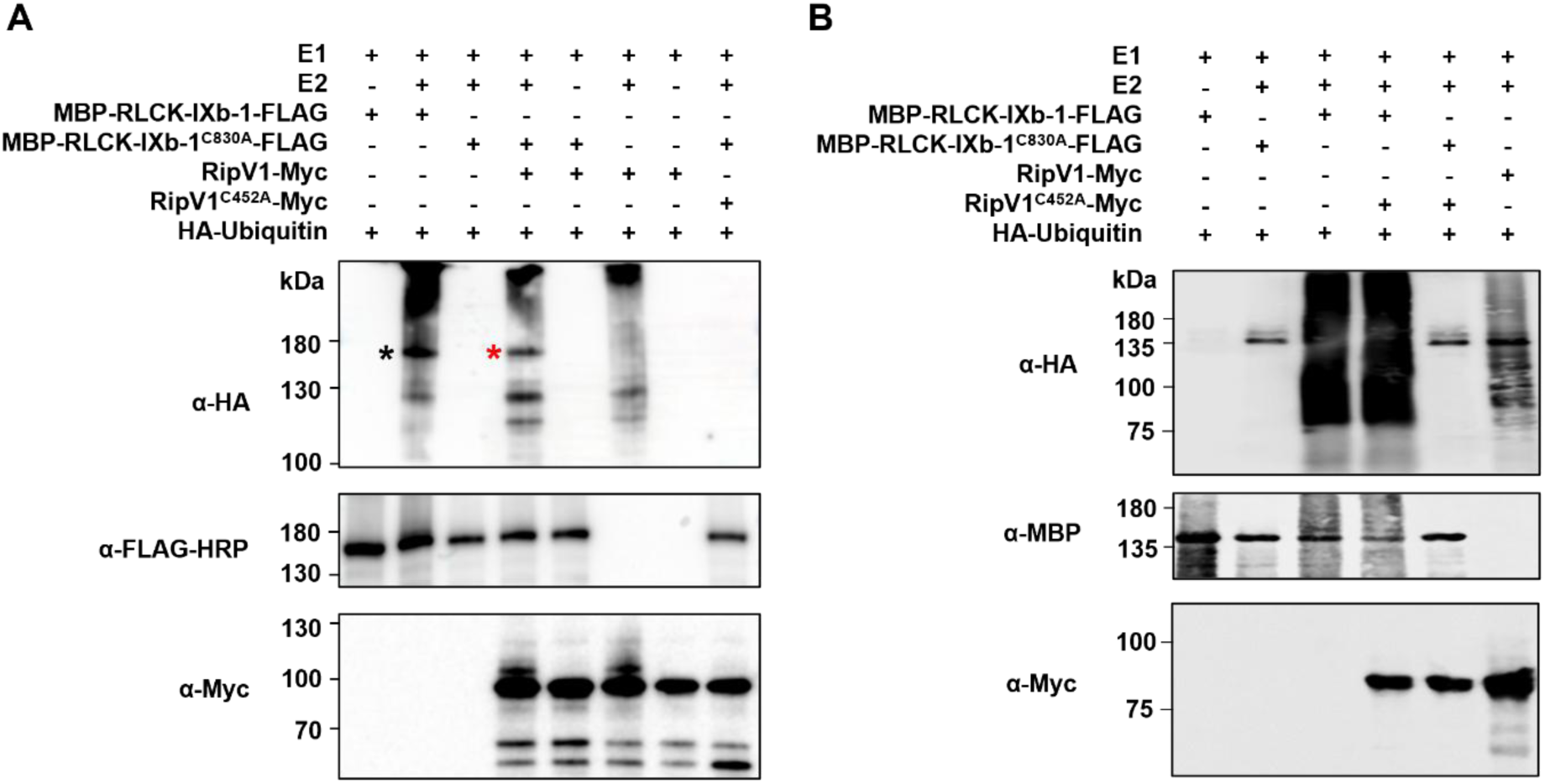
RipV1 ubiquitinates RLCK-IXb-1 *in vitro*. (**A**) E3 ligase-dead mutant (RLCK-IXb-1^C830A^) was ubiquitinated in the presence of RipV1 *in vitro*. FLAG-tagged RLCK-IXb-1 and E3 ligase-dead mutant (RLCK-IXb-1^C830A^) and Myc-tagged RipV1 and E3 ligase-dead mutant (RipV1^C452A^) recombinant proteins were incubated with HA-Ubiquitin in the presence or absence of human E1 (UBE1) and E2 (UbcH5B). Reaction mixtures were separated by SDS-PAGE and detected by immunoblot assay using anti-HA, anti-FLAG-HRP and anti-Myc antibodies. The black asterisk marks a band near the expected size of recombinant RLCK-IXb-1 (146 kDa). Red asterisk indicates ubiquitinated RLCK-IXb-1^C830A^ in presence of RipV1. (**B**) Ubiquitination of RipV1 by RLCK-IXb-1 was undetectable *in vitro*. Reaction mixtures were separated by SDS-PAGE and detected by immunoblot assay using anti-HA, anti-MBP and anti-Myc antibodies.

We next compared ubiquitination patterns in reactions containing either RLCK-IXb-1, RLCK-IXb-1^C380A^, or RipV1 alone, or RLCK-IXb-1^C380A^ and RipV1 together (Figure 4A). While no ubiquitination signal was observed on RLCK-IXb-1^C380A^ alone, we observed a ubiquitination pattern on RLCK-IXb-1^C380A^ when co-incubated with RipV1, including a band corresponding to RLCK-IXb-1 protein size (Figure 4A, red asterisk). A similar band was detected in the RLCK-IXb-1 auto-ubiquitination reaction (Figure 4A, black asterisk) but was not observed in the RipV1 auto- ubiquitination reaction. These results indicate that RipV1 could trans-ubiquitinate RLCK-IXb-1, possibly on residues that participate in the auto-ubiquitination of RLCK-IXb-1. Additionally, we attempted to test if RLCK-IXb-1 could trans-ubiquitinate RipV1 by comparing ubiquitination patterns in reactions containing RLCK-IXb-1 or RipV1 alone; RipV1^C452A^ and RLCK-IXb-1 together; and RipV1^C452A^ and RLCK-IXb-1^C380A^ together (Figure 4B). Although the reaction containing RLCK-IXb-1 and RipV1^C452A^ produced an intense ubiquitination signal compared to the reaction containing only RipV1^C452A^, this was undistinguishable from the intense signal produced by RLCK-IXb-1 auto-ubiquitination (Figure 4B). RipV1 detection with anti-Myc antibodies revealed a faint higher molecular weight signal for RipV1 but not for RipV1^C452A^ (Figure 4). As this signal was not observed in the sample containing RipV1^C452A^ and RLCK-IXb-1 (Figure 4B), these data suggest that RipV1 is unlikely to be ubiquitinated by RLCK-IXb-1.

Ubiquitination can have a major impact on protein stability (Sadanandom et al., 2012; Zhang et al., 2015). As RipV1 could ubiquitinate RLCK-IXb-1 *in vitro*, we next investigated whether RipV1 could affect RLCK-IXb-1 stability using transient expression in *N. benthamiana*. RLCK-IXb-1-HA was co-expressed with GFP, RipV1-YFP, or RipV1^C452A^-YFP and protein accumulation was detected by immunoblot (Figure 5). RLCK-IXb-1 accumulation was the highest in the presence of RipV1 in different assays (Figures 2A and 5A). To compare relative protein abundance, the band intensity of RLCK-IXb-1 detected with anti-HA antibodies were quantified across multiple independent experiments. RLCK-IXb-1 accumulation was the highest in the presence of RipV1 compared to the catalytically dead RipV1^C452A^ (Figure 5B). These results suggest that RLCK-IXb-1 is stabilized in the presence of RipV1 with a functional E3 ligase activity. Taken together with our finding that RipV1 can ubiquitinate RLCK-IXb-1, this experiment indicates that RipV1-mediated ubiquitination enhances RLCK-IXb-1 stability *in planta*.

**Figure 5.**
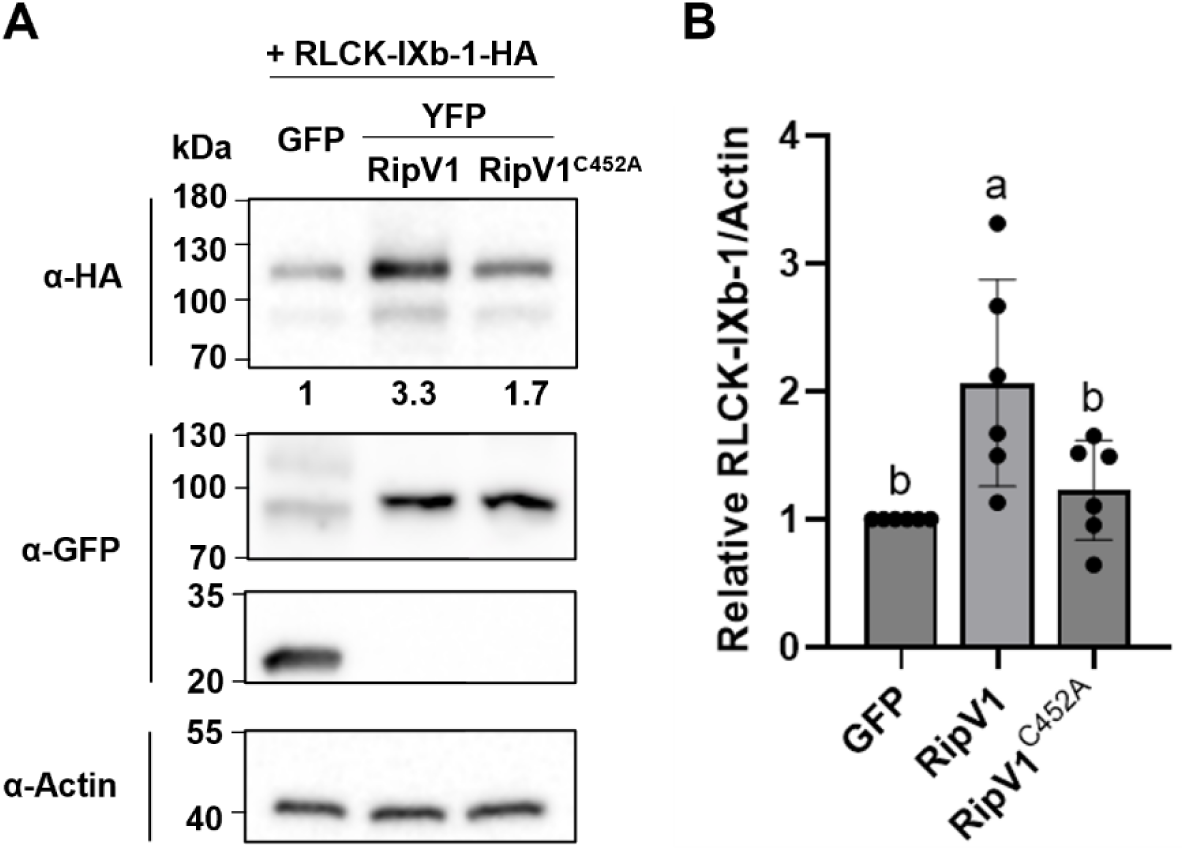
RLCK-IXb-1 stability is promoted in the presence of RipV1 *in planta*. (**A**) RLCK-IXb- 1-HA was transiently co-expressed with GFP, RipV1-YFP or E3 ligase-dead RipV1^C452A^-YFP in *N. benthamiana*. Leaf samples were harvested 45 hours post infiltration. Total protein extracts were probed with anti-HA, anti-GFP and anti-Actin antibodies. Accumulation of the indicated proteins was measured using ImageJ. Numbers under the anti-HA antibody blot indicate relative protein accumulation normalized by actin. (**B**) Quantification of RLCK-IXb-1 accumulation in the presence of GFP, RipV1 or RipV1^C452A^. Relative differences in RLCK-IXb-1 accumulation were quantified in six independent experiments, including the one shown in (A). Data are presented as the mean normalized band intensity values ± standard deviation. Different letters indicate statistically significant differences analyzed by one-way analysis of variance followed by Tukey’s multiple comparisons test (*P* < 0.01).

### RLCK-IXb-1/2 act as negative regulators of plant defense responses

The phylogenetic analysis of the RLCK-IXb subfamily in Arabidopsis, tomato, and *N. benthamiana* identified the two closest *N. benthamiana* homologs for each of the RLCK-IXb proteins that interact with RipV1 (Figure S2). Evolutionary comparison revealed that RLCK-IXb-1 and -2 were the most closely related and clustered together, implying potential functional similarity. To test if NbRLCK-IXb proteins play a role in plant immune responses, we silenced either *NbRLCK-IXb-1*, *-2*, *-3* and *-9* together or *NbRLCK-IXb-1* and *-2* together to account for potential redundancy. The efficiency of silencing was confirmed by qRT-PCR (Figure S7). The ROS production elicited by flg22 treatment was significantly increased in *N. benthamiana* plants silenced for *NbRLCK-IXb-1/2/3/9* compared to the control silenced plants (Figure 6A). This could be attributed to *NbRLCK-IXb-1* and *-2*, as silencing only *NbRLCK-IXb-1/2* also resulted in enhanced ROS production upon flg22 treatment (Figure 6B). These results suggest that at least two members of the RLCK-IXb family that interact with RipV1 function as negative regulators of the early immune response in *N. benthamiana*, as also suggested by their close evolutionary relationship.

**Figure 6.**
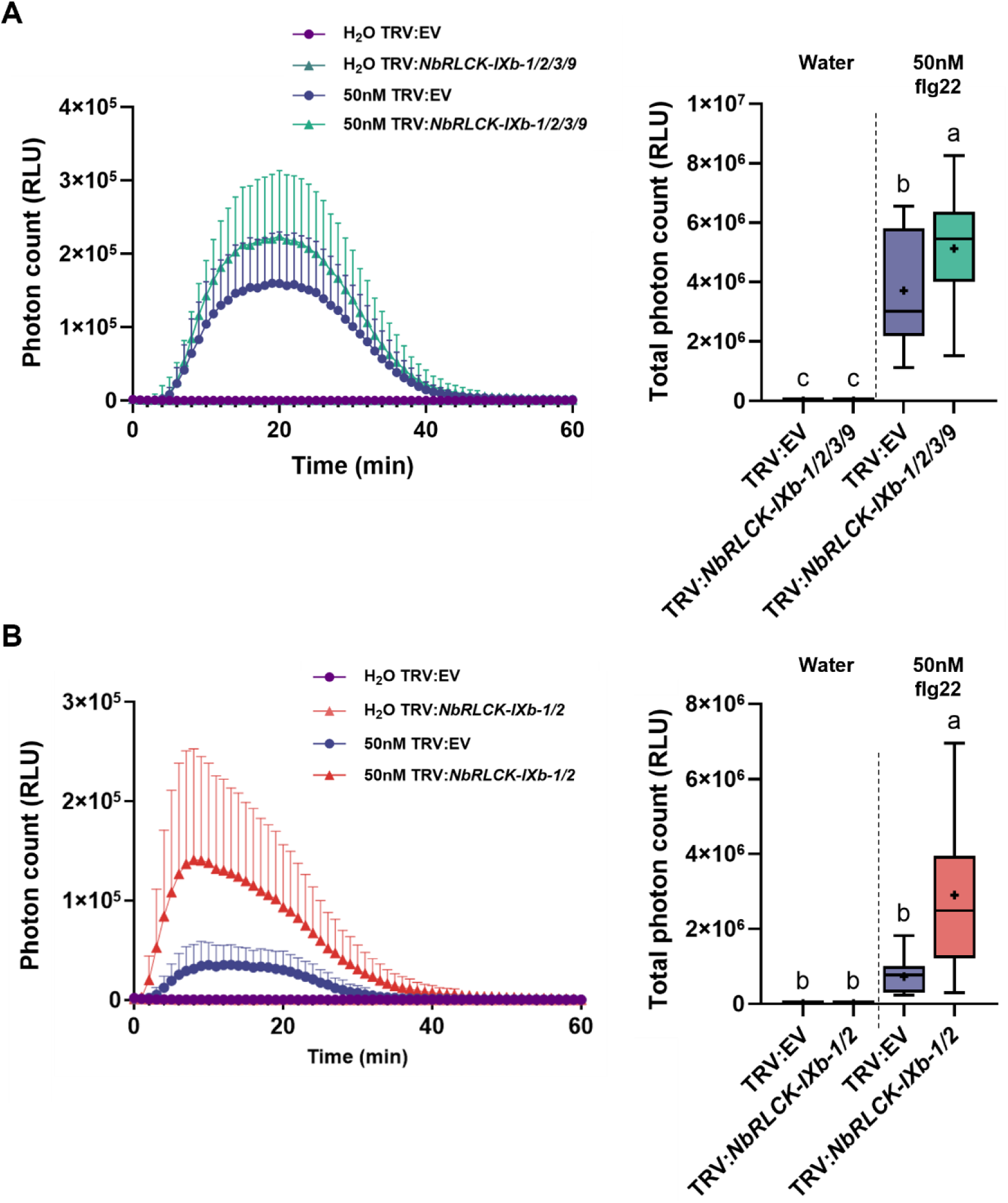
RLCK-IXb-1/2 act as negative regulators of flg22-induced ROS in *N. benthamiana*. The immunogenic flagellin peptide (flg22)-triggered ROS production was elevated in TRV:*NbRLCK-IXb-1/2/3/9* plants (**A**) and TRV:*NbRLCK-IXb-1/2* plants **(B)**. *N. benthamiana* plants silenced for *NbRLCK-IXb-1*, *-2*, *-3*, and *-9* (TRV:*NbRLCK-IXb-1/2/3/9)*, *NbRLCK-IXb-1* and *-2* (TRV:*NbRLCK-IXb-1/2*) or empty vector (TRV:EV) were compared. Leaf disks were treated with water or 50 nM flg22 and ROS production was monitored for 60 minutes. Kinetic curves present the mean values ± standard deviation of relative light units (RLU) from a representative experiment out of four (A) and three (B) independent experiments (n = 64 in A; n = 48 in B). Box- and-whisker plots on the right show total ROS production. Boxes indicate interquartile range, central lines indicate the median, whiskers indicate the minimum and maximum values and ‘+’ indicates the mean. Different letters denote statistically significant differences analyzed by one- way analysis of variance followed by Tukey’s multiple comparisons test (*P* < 0.0001).

### RLCK-IXb-1 negatively regulates RipV1-induced cell death

RipV1 contributes to *R. solanacearum* virulence on potato and induces cell death in an E3 ligase -dependent manner in *N. benthamiana* (Cheng et al., 2024). To investigate whether RipV1- induced cell death is the consequence of activated NLR signaling or results from RipV1 virulence activity, we observed RipV1-induced cell death in *N. benthamiana* plants silenced for *NbSGT1*, a central regulator of immune receptor function (Austin et al., 2002; Azevedo et al., 2002). The *R. solanacearum* effectors RipA1 and RipE1 were used as controls for SGT1-independent and SGT1- dependent cell death, respectively (Jeon et al., 2020; Landry et al., 2020; Sang et al., 2020; Solé et al., 2012). In plants silenced for *NbSGT1* using tobacco rattle virus (TRV)*:NbSGT1*, RipV1- induced cell death was delayed compared to the robust cell death observed at 36 hours post inoculation (hpi) in control plants transformed with the empty vector (TRV:EV) (Figure S8A and S8B). Because *NbSGT1* silencing is known to also affect protein stability (Yu et al., 2019), we assessed RipV1 protein accumulation in the silenced plants by immunoblot. Similar to FLAG-GFP accumulation, RipV1 protein level was reduced in TRV*:NbSGT1* tissues (Figure S8C), possibly explaining the delayed cell death. We further quantified RipV1-induced cell death in *N. benthamiana* knock-out lines lacking essential components of immune receptor signaling: the *EDS1/PAD4/SAG101* module (*epss*; Lapin et al., 2019), *NRG1* (*nrg1*; Qi et al., 2018) or *NLR REQUIRED FOR CELL DEATH 2/3/4* (*nrc2/3/4*; Wu et al., 2020) (Figure S8D and S8E). RipV1- induced cell death was similar in the wild-type, *epss*, *nrg1* and *nrc2/3/4* lines, suggesting that the NLR signaling network is dispensable for this response.

Next, we investigated whether RipV1-interacting RLCKs could regulate RipV1-induced cell death. In TRV*:NbRLCK-IXb-1/2/3/9* plants, but not in TRV:EV control plants, RipV1-induced cell death was detectable when RipV1 was expressed at a low level (Figure 7A and 7B). Conversely, RipV1-induced cell death was reduced in leaf tissues where RLCK-IXb-1 and RipV1 were co-expressed (Figure 7C and D). We confirmed that RipV1 accumulates to a comparable level in TRV:EV and TRV*:NbRLCK-IXb-1/2/3/9* silenced plants (Figure S9A) and that both RipV1 and RLCK proteins were detectable in the co-expression assay (Figure S9B). Taken together, these findings support the role of RLCK-IXb-1 as a negative regulator of RipV1-induced cell death.

**Figure 7.**
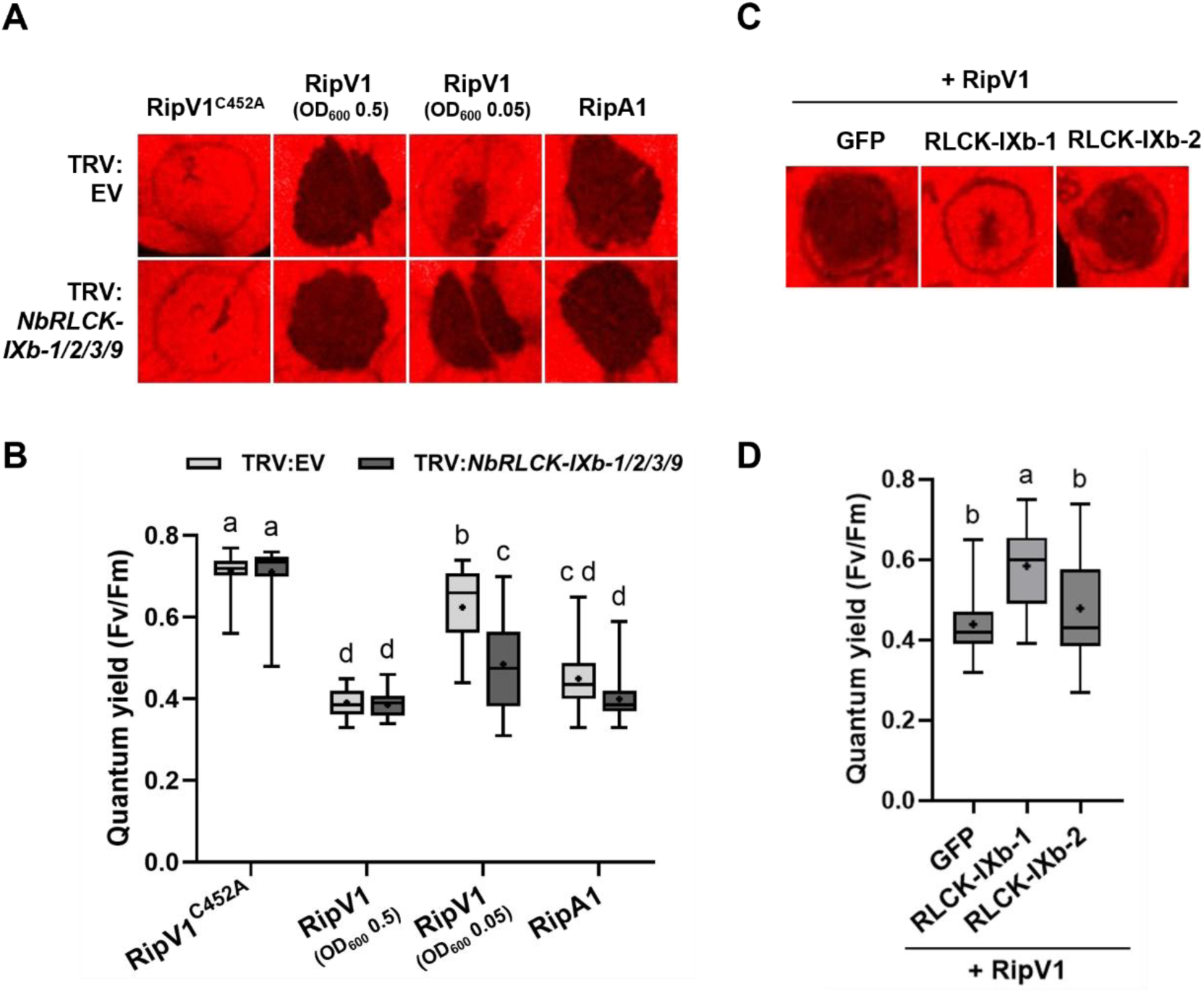
RLCK-IXb-1 negatively regulates RipV1-induced cell death in *N. benthamiana*. (**A**) Cell death triggered by RipV1 was facilitated in TRV:*NbRLCK-IXb-1/2/3/9* plants. RipV1^C452A^-FLAG, RipV1-FLAG and RipA1-FLAG were transiently expressed by agroinfiltration in TRV:EV and TRV:*NbRLCK-IXb-1/2/3/9* plants. Leaves were photographed 3 days post infiltration (dpi). False- color images representing quantum yield of chlorophyll (QY) were generated using FluorCam software. A customized color scale was applied to enhance visual contrast, in which black regions indicate areas of cell death. (**B**) Box-and-whisker plot shows QY values from three independent repeats (n = 16) of the experiment shown in (A). Boxes indicate interquartile range, the central line indicates the median, whiskers indicate the minimum and maximum values and ‘+’ indicates the mean. Different letters denote statistically significant differences as analyzed by two-way analysis of variance (ANOVA) followed by Tukey’s multiple comparison test (*P* < 0.0001). (**C**) Expression of RLCK-IXb-1 delayed RipV1-induced cell death in *N. benthamiana*. RipV1-FLAG was co-expressed with GFP, RLCK-IXb-1-HA or RLCK-IXb-2-HA. Leaves were photographed 3 dpi. (**D**) Box-and-whisker plot shows QY values from 4 independent repeats (n = 33) of the experiment shown in (C). Boxes indicate interquartile range, the central line indicates the median, whiskers indicate the minimum and maximum values and ‘+’ indicates the mean. Different letters denote statistically significant differences as analyzed by one-way ANOVA followed by Tukey’s multiple comparison test (*P* < 0.0001).

## DISCUSSION

RLCKs comprise multiple subfamilies and some contribute redundantly to plant immune responses (Huang et al., 2024; Jalilian et al., 2023; Lin et al., 2013; Shiu et al., 2004). Several members of the RLCK-VII subfamily in Arabidopsis show functional redundancy in flg22-induced immune signaling (Bi et al., 2018; Rao et al., 2018). Additionally, different classes of RLCK-VII proteins regulate ROS production and cell death elicited by *Phytophthora palmitovora* in *N. benthamiana* (Huang et al., 2024). Here, we identified four tomato RLCKs in subfamily IXb as RipV1 interactors. Because genetics is challenging in tomato, we opted to test the functional role of these RLCKs in the close relative *N. benthamiana*, which has well-developed genetics methods. Silencing the *N. benthamiana* RLCK-IXb homologs enhanced flg22-triggered ROS burst, suggesting that at least RLCK-IXb-1 and -2 act as redundant negative regulators of early immune responses. Whether this function relies on their catalytic activity remains to be evaluated. RipV1 interacting-RLCKs harbor both kinase and PUB domains. While RLCK-IXb-1 displayed E3 ligase activity, no kinase activity was detected in our experimental conditions, suggesting that it may be an inactive kinase or that additional cellular cues are required for its activation.

Phylogenetic analysis revealed that tomato RLCK-IXb-1 clusters most closely with AtPUB33, a characterized member of PUB class VI (Dong et al., 2023; McLellan et al., 2022; Trenner et al., 2022). Arabidopsis *pub33* mutants are impaired in flg22-triggered ROS production and defense gene expression, indicating a positive role in plant immune signaling (Dong et al., 2023). StUBK and NbUBK, which have previously been reported as homologs of AtPUB33 in potato and *N. benthamiana*, are included in the RLCK-IXb-1 cluster and positively regulate immune responses to *P. infestans* (He et al., 2019; McLellan et al., 2022). However, NbPUB33 referred as a *N. benthamiana* homolog of AtPUB33 (Dong et al., 2023) is more closely related to tomato RLCK-IXb-13 (Figure S2). Thus, a reevaluation of orthologous relationship within the AtPUB33 clade appears necessary. Nonetheless, our results further support RLCK-IXb subfamily members as regulators of plant immune signaling. Moreover, StUBK and AtPUB33 were identified as interactors of the *P. infestans* effector Pi06087 and the *Apolygus lucorum* effector AI106, respectively (Dong et al., 2023; He et al., 2019; McLellan et al., 2022). Together with our finding that RipV1 targets RLCK-IXb-1, this suggests that proteins within the AtPUB33 clade may serve as common targets for pathogen effectors and could be leveraged to develop disease resistance strategies (McLellan et al., 2022).

Different types of ubiquitin linkages determine distinct cellular outcomes for their protein targets (Zhou and Zeng, 2017). BIK1 undergoes polyubiquitination by PUB25/26 promoting degradation, while monoubiquitination by RHA3A/B leads to BIK1 redistribution (Ma et al., 2020;

Wang et al., 2018). Monoubiquitination has been reported to inhibit polyubiquitination via structural mechanisms or lysine residue specificity (de Bie and Ciechanover, 2011; Herrador et al., 2013; Sadowski et al., 2012). Our observations indicate that RLCK-IXb-1 can auto-ubiquitinate and be trans-ubiquitinated by RipV1, possibly on similar residues. RLCK-IXb-1 may regulate its own stability through auto-ubiquitination, a common property of PUBs (Trenner et al., 2022). The increased stability of RLCK-IXb-1 in the presence of RipV1 could be due to the ubiquitination mediated by RipV1. Mapping auto- and trans-ubiquitination sites on RLCK-1Xb would provide deeper understanding of the biochemical interaction between RipV1 and RLCK- IX-1.

It remains to be clarified whether the delayed RipV1-induced cell death upon co-expression with RLCK-IXb-1 results from RLCK-IXb-1 stabilization. A similar regulatory strategy has been described for *P. infestans* effector AVR3a, which stabilizes the host E3 ligase CMPG1 to suppress INF1-triggered cell death (Bos et al., 2010). Although in this case the effector modulates a distinct immune input, it illustrates how effectors can stabilize host proteins that modulate cell death timing or intensity. RipV1-induced cell death in *N. benthamiana* depends on its E3 ligase activity (Cheng et al., 2024). Whether this cell death represents a case of effector recognition by immune receptors was not yet addressed. Although *NbSGT1* silencing delayed RipV1-induced cell death, RipV1 protein accumulation was reduced in TRV:*NbSGT1* plants, potentially due to a stabilizing role of SGT1, as observed for the *Salmonella* NEL effector SspH2 (Bhavsar et al., 2013). Additionally, we evaluated RipV1-triggered cell death in knockout lines lacking canonical NLR signaling factors. RipV1 retained its ability to induce cell death, indicating that this response does not require helper NLRs or the EDS1/PAD4/SAG101 complex. A similar situation is observed for RipAW, another NEL effector that elicits *EDS1*- and *NRG1*-independent cell death in *N. benthamiana* (Ouyang et al., 2023). Collectively, these results suggest that RipV1- or other NEL effector-induced cell death might depend on singleton NLRs that mediate immune signaling independently (Contreras et al., 2023). Given that NEL effectors contribute to the virulence of *R. solanacearum*, a deeper characterization of the functional role of RLCK-IXb-1 in immune signaling could offer new insights into the mechanisms underlying *R. solanacearum* virulence in host plants.

## EXPERIMENTAL PROCEDURES

### Plant growth conditions and bacterial strains

*A. N. benthamiana* plants were grown at 24 °C under 16 h light/8 h dark photoperiod for 4-5 weeks in a controlled plant growth room. The *N. benthamiana* knock-out lines used in this study were previously described: *epss* (Lapin et al., 2019), *nrg1* (Qi et al., 2018), and *nrc2/3/4* (Wu et al., 2020). *Escherichia coli* DH5α and Rosetta (DE3), and *Agrobacterium tumefaciens* AGL1 were grown at 37 °C and 28 °C respectively on Lysogeny Broth (LB) medium supplemented with appropriate antibiotics.

### Molecular constructs

Modules for *R. solanacearum* effectors compatible for Golden Gate assembly were previously generated and described in Prokchorchik et al. (2020). Coding sequences of tomato RLCK genes were obtained from Sakamoto et al. (2012) and divided into modules of approximately 1 kb in size with flanking BsaI sites for Golden Gate assembly (Engler et al., 2008). Modules were amplified by PCR from tomato Heinz cDNA or synthesized (Cosmogenetech) and subsequently cloned into pICH41021. Site-directed mutagenesis was performed by whole-plasmid PCR using High-Fidelity DNA polymerase (Thermo) and complementary mutagenic primers (Table S2) containing the desired nucleotide substitutions. Following amplification, PCR products were treated with DpnI (NEB) to digest the parental methylated plasmid DNA. The resulting plasmids were transformed into *E. coli* DH5α and all mutations were confirmed by Sanger sequencing (Macrogen).

### Yeast two-hybrid assay

Yeast strain EGY48 carrying the lacZ-bearing reporter plasmid pSH18-34 was transformed with pLexA constructs containing *R. solanacearum* effectors. Transformants were selected on synthetic defined (SD) minimal agar media lacking uracil and histidine (Clontech) for 2-3 days. Similarly, yeast strain RFY206 was transformed with pB42-AD constructs containing tomato RLCKs and transformants were selected on SD media lacking tryptophan (Clontech). Plasmid DNA was extracted from overnight yeast culture using plasmid extraction kit (QIAGEN). The presence of coding sequence was confirmed by amplification using primers (Table S2) from each vector. Transformants were grown overnight in 200 μl of yeast extract peptone dextrose (YPD) broth for mating. Overnight cultures were grown on SD media lacking histidine, uracil, and tryptophan and were incubated at 30 °C. Mated cells were spotted on SD media lacking histidine, uracil, leucine, and tryptophan (Clontech) containing raffinose, galactose, and X-gal (Sigma) and grown for 2 to 5 days before being photographed.

### Agrobacterium-mediated transient expression

Binary constructs (pICH86988) were introduced into *A. tumefaciens* strain AGL1 through electroporation. Transformants were grown on LB agar medium containing selective antibiotics for 2-3 days. Single colonies were inoculated into LB liquid media with selective antibiotics and cultured overnight. Overnight cultures were centrifuged at 5,400 x *g* for 3 minutes and resuspended in agroinfiltration buffer (10 mM MgCl_2_ and 10 mM MES-KOH, pH 5.6). Cell suspensions OD_600_ were adjusted depending on experiments. For cell death observation, *A. tumefaciens* cultures carrying RLCK-IXb-1-HA or RLCK-IXb-2-HA were adjusted to an OD_600_ of 1.0, while all other cultures were prepared at an OD_600_ of 0.5 unless otherwise noted. For protein accumulation assays, *A. tumefaciens* cultures carrying RipV1-YFP, GFP, and RLCK-IXb-1-HA were adjusted to an OD_600_ values of 0.1, 0.1, and 0.2, respectively. For confocal microscopy, *A. tumefaciens* cultures carrying RipV1-YFP, RLCK-IXb-1-YFP or CFP, and AtFLS2-mCherry were adjusted to OD_600_ values of 0.1, 0.4, and 0.4, respectively. In all experiments involving RLCK-IXb- 1-HA or RLCK-IXb-2-HA, *A. tumefaciens* cultures carrying P19 (Lakatos et al., 2004) was added at OD_600_ of 0.1. When multiple suspensions were co-infiltrated, they were mixed in equal volumes. The final suspensions were infiltrated into 5-week-old *N. benthamiana* leaves using a needleless syringe.

### Protein extraction and immunoprecipitation

*A. N. benthamiana* agroinfiltrated leaf samples were ground to a fine powder in liquid nitrogen. Total proteins were extracted in GTEN extraction buffer (10 % glycerol, 50 mM Tris-HCl pH 7.5, 2 mM EDTA pH 8, 150 mM NaCl) including 5 mM Dithiothreitol (DTT), 0.2 % Nonidet P (NP)-40 (Sigma-Aldrich), 1 % Polyvinylpolypyrrolidone (PVPP), 1 tablet of cOmplete protease inhibitor cocktail per 40 ml (Roche). Extracts were cleared by centrifugation at 15,000 x *g* for 10 minutes at 4 °C and filtered through MiraCloth (Millipore). The filtered proteins were denatured in 3X Sodium dodecyl sulfate (SDS) sample buffer at 96 °C for 10 minutes. For immunoprecipitation, 1.5 ml of total protein extracts were incubated with 30 μl of GFP-Trap agarose beads (ChromoTek) for 2 hours at 4 °C. The beads were washed three times with extraction buffer without PVPP and denatured in 50 μl of 3x SDS sample buffer at 96 °C for 10 minutes. The proteins were separated by SDS-polyacrylamide gel electrophoresis (PAGE) and probed with anti-HA (Santa Cruz), anti-GFP (Santa Cruz), FLAG-HRP (Sigma-Aldrich) or anti-Actin (Agrisera) antibodies. Detection was performed using SuperSignalTM West Substrate (Thermo Scientific) and visualized with an Azure 600 (Azure Biosystems).

### Confocal microscopy

Confocal imaging was performed using a Leica SP8X confocal laser scanning microscope equipped with a 40x water immersion objective lens. Fluorescence was observed at 40-45 hpi in leaves expressing RipV1-YFP, and at 48 hpi for all other samples. YFP, CFP, and mCherry were excited at 514 nm (argon laser), 405 nm (diode laser), and 594 nm (helium-neon laser), respectively. Emission was collected at 525-565 nm (YFP), 470-502 nm (CFP), and 609-662 nm (mCherry). Images were processed using LAS X software (Leica Microsystems).

### Recombinant protein purification

RipV1 and RLCK-IXb-1 constructs in pOPIN vectors (Choi et al., 2021) with C-terminal Myc and FLAG epitope tags were transformed into *E. coli* Rosetta (DE3) for protein expression. Rosetta cells carrying constructs were grown at 37 °C. Protein expression was induced by adding 1 mM of isopropyl β-D-1-thiogalactopyranoside (IPTG) when OD_600_ value reached 0.8. Following IPTG induction, RipV1 and RLCK-IXb-1 were cultured at 37 °C and 28 °C for 16 hours respectively. Cells were centrifuged at ∼1900 x *g* for 15 minutes at 4 °C and resuspended with wash buffer (50 mM Tris-HCl pH 8.0, 300 mM NaCl, 10 mM Imidazole, 0.1 mM Ethylenediaminetetraacetic acid (EDTA), 1 mM Phenylmethylsulfonyl fluoride (PMSF). Resuspended cells were sonicated and centrifuged at 12,000 x *g* for 10 minutes at 4 °C. Supernatants were filtered using 0.45 µm filter and loaded onto a column containing HisPur^TM^ Cobalt Resin (Thermo). Samples were washed two times with wash buffer and eluted with elution buffer (50 mM Tris-Cl pH 8.0, 50 mM NaCl, 300 mM Imidazole, 0.1 mM EDTA, 1 mM PMSF). Protein concentration of recombinant samples was determined using Pierce™ Bradford Plus Protein Assay Reagent (Thermo) according to the manufacturer’s instructions. Protein concentrations were determined by measuring absorbance at 595 nm and calculated based on the linear range of the bovine serum albumine standard curve.

### *In vitro* ubiquitination assay

Ubiquitination assays were conducted following the method from Nakano et al. (2017) with some modifications. In the reaction mixture, the amount of UBE1 and UbcH5B (Boston Biochem) were changed to 0.05 µg and 1 µg, respectively. For auto-ubiquitination assays of RipV1 or RLCK-IXb- 1, 1 µg of each purified protein was used. For trans-ubiquitination assays, 1 µg RipV1 and 3 µg RLCK-IXb-1 of purified proteins were used. Reactions were incubated at 37 °C for 1 to 3 hours. Reactions were stopped by adding 2x Laemmli buffer and heating at 96 °C for 10 min. Ubiquitination assays were analyzed by SDS-PAGE followed by immunoblot assay with anti-HA (Santa Cruz), anti-Myc (Cell Signaling Technology), and anti-FLAG-HRP (Sigma-Aldrich) antibodies. Detection was performed using SuperSignal^TM^ West Substrate (Thermo Scientific) and visualized with an Azure 600 (Azure Biosystems).

### *In vitro* kinase assay

*In vitro* kinase assays were conducted using 2 μg kinase in a buffer containing 50 mM Tris-HCl pH 8.0, 25 mM MgCl_2_ and 25 mM MnCl_2_, 5 mM DTT, 5 μM ATP, and 0.5 μCi γ^32^P-ATP. All reactions were incubated at 30°C for 45 minutes. Reactions were stopped by adding 5x Laemmli buffer and heating at 80 °C for 5 min. Proteins were separated on 8% SDS-PAGE gels, run at 80 V for 30 min, followed by 150 V for 1 h. The gels were sandwiched between two sheets of transparency film, exposed to a storage phosphor screen (Cytiva) overnight and visualized using Typhoon FLA 9500 imager (Cytiva/Amersham) or a Sapphire FL Phosphorimager (Azure Biosystems). Gels were then stained with Coomassie Brilliant Blue (CBB) R-250 (MP Biomedicals) and scanned (HP Officejet Pro 8620) for total protein visualization.

### Virus-induced gene silencing

*A. tumefaciens* carrying pTRV1 and pTRV2 (Ahn et al., 2023; Choi et al., 2021; Liu et al., 2002) at OD_600_ of 0.5 were co-infiltrated into 2-week-old *N. benthamiana* leaves. *N. benthamiana* plants were further grown for 4-5 weeks before use in experiments. To design specific silencing fragments, coding sequences of tomato RLCKs homologs in *N. benthamiana* were analyzed with the SolGenomics Network VIGS tool (http://vigs.solgenomics.net). The following *N. benthamiana* gene models were targeted for silencing: *NbRLCK-IXb-1-1* (*Niben261Chr03g0421011.1*), *NbRLCK-IXb-1-2* (*Niben261Chr09g1137026.1*), *NbRLCK-IXb-2-1* (*Niben261Chr06g0821015.1*), *NbRLCK-IXb-2-2* (*Niben261Chr08g0471021.1*), *NbRLCK-IXb-3-1* (*Niben261Chr06g0936005.1*), *NbRLCK-IXb-3-2* (*Niben261Chr08g0547001.1*), *NbRLCK-IXb-9-1* (*Niben261Chr12g0682001.1*), and *NbRLCK-IXb-9-2* (*Niben261Chr19g0053013.1*). Fragments (∼150 bp) were amplified with overhangs compatible for Golden Gate Assembly and assembled in pTRV2 vector.

### Measurement of ROS production

Leaf disks were collected from 5-week-old silenced plants and incubated in 150 μl of distilled water overnight. The distilled water was replaced by 100 μl of reaction solution containing 100 μM luminol (Sigma-Aldrich), 2 μg horseradish peroxidase (Sigma-Aldrich) and 50 nM flg22 (Peptron) or the equivalent volume of water. Luminescence was measured every minute for 60 minutes using a SYNERGY HTX multimode microplate reader (BioTek) and expressed as relative light units (RLU).

### Cell death quantification

Cell death intensity was evaluated by measuring the maximum quantum yield of chlorophyll in photosystem II (F*v*/F*m*), as an indicator of photosynthetic efficiency (Lee et al., 2021). Agroinfiltrated leaves were collected at the indicated time point and immediately transferred into the closed imaging chamber. Chlorophyll fluorescence was captured using standard FluorCam settings (Photon Systems Instruments).

### Protein domain and 3D structure prediction

Protein domain prediction was conducted using NCBI Conserved Domain (CD) search tool (https://www.ncbi.nlm.nih.gov/Structure/cdd/wrpsb.cgi). Protein structure prediction was performed using AlphaFold 3 (Abramson et al., 2024; AlphaFold Server, https://alphafoldserver.com/). Structure prediction was conducted using default parameters for monomer mode. 3D structures were visualized and annotated in *CCP*4*mg* molecular-graphics software (McNicholas et al., 2011).

### Multiple sequence alignment and phylogenetic analysis

Amino acid sequences were aligned using ClustalW (Thompson et al., 1994; EMBL-EBI web server, https://www.ebi.ac.uk/Tools/msa/clustalw2/). The resulting alignment was visualized using ESPript 3.0 (Robert and Gouet, 2014; https://espript.ibcp.fr/). Phylogenetic analysis was conducted using Geneious Prime (Dotmatics; version 2025.1.1). Amino acid sequences were aligned using the global alignment algorithm with the BLOSUM62 substitution matrix. A phylogenetic tree was constructed using neighbor-joining method based on the Jukes-Cantor model.

### Quantitative real-time PCR (qRT-PCR)

Total RNA was extracted from leaves of *NbRLCK-IXb-1/2/3/9*-silenced and control plants using TRIzol reagent (Invitrogen). cDNA was prepared from 2 µg of RNA using Maxima First Strand cDNA Synthesis Kit for RT-qPCR (Thermo). qPCR reactions were performed using GoTaq® qPCR Master Mix (Promega), following the manufacturer’s instructions. Expression levels were normalized to *NbEF1α* (Table S2) as the reference gene using the ΔΔ^Ct^ method (Livak and Schmittgen, 2001).

### Statistical analysis

All the experiments included technical replicates and were performed at least two or three times independently, as indicated in figure legends. Unless otherwise stated, values from independent biological repeats were merged for statistical analyses. Statistical significance of differences was analyzed by analysis of variance (ANOVA) and post-test as indicated in the figure legends using GraphPad Prism (version 10.4.2).

## Supporting information

Choi J et al_supp info

## ACKNOWLEDGMENTS

This work was supported by the National Research Foundation of Korea (NRF) funded by the Korean Ministry of Sciences and ICT (Projects No RS-2018-NR031006 and No RS-2024-00349151) to CS, as well as the Natural Sciences and Engineering Research Council of Canada (NSERC) Discovery Program (RGPIN-2016-04787; RGPAS-492902-2016; RGPIN-2024-04072) to JM. The authors declare no conflict of interest.

---

**Figure S1.** 3D structure models of RipV1 and RLCK-IXb-1. (A, **B)** RipV1 protein structure with predicted local distance difference test (plDDT) confidence score (A) and colored domains (B). RipV1 truncation constructs used in yeast two-hybrid assays are shown in (B). The grey region (1- 374 aa) corresponds to the N-terminal domain, and the navy region (374-685 aa) includes the NEL domain (374-604 aa; conserved domain database (CDD) cl38455). **(C, D)** RLCK-IXb-1 protein structure with plDDT confidence score (C) and colored regions (D). RLCK-IXb-1 domains were defined according to the domain analysis of RLCK-IXb-1: USP (1-208 aa; brown; cd01989), Smc (361-520 aa; maroon; cl34310), Kinase (521-814 aa; dark magenta; cl21453), and PUB (815-894 aa; violet; cd16655) domains. The unfolded region (209-360 aa) between USP and Smc domains is colored in grey. Domain analysis was conducted by NCBI CD-search and 3D structure was predicted by AlphaFold 3.

**Figure S2.** Phylogenetic analysis of RLCK subfamily IXb in Arabidopsis, tomato, and *N. benthamiana*. Arabidopsis and tomato RLCK-IXb protein sequences were obtained from Sakamoto et al. (2012). RLCK-IXb members were retrieved from the *N. benthamiana* genome (v2.6.1) available at (https://solgenomics.net/organism/Nicotiana_benthamiana/genome) using the 18 Arabidopsis and 15 tomato RLCK-IXb protein sequences as queries. The RLCK-XII XOPJ4 IMMUNITY 2 (JIM2) was used as an outgroup (Schultink et al., 2019). The tree was built on full-length amino acid sequences using the neighbor-joining method with 1,000 bootstrap replicates performed by the Jukes-Cantor model. The scale bar indicates evolutionary distance. RLCKs interacting with RipV1 in yeast two-hybrid are highlighted in blue. *N. benthamiana* homologs of RLCK-IXb-1, -2, -3, and -9 targeted by virus-induced gene silencing are highlighted in green. The protein highlighted in magenta is identical to NbD000250.1 reported as AtPUB33 homolog by Dong et al. (2023). Another AtPUB33 homolog, NbUBK, described in He et al. (2018) is labeled in purple. Members of RLCK subfamily IXb that lack a PUB domain are shown in italics.

**Figure S3.** Subcellular localization of RipV1 and RLCK-IXb-1. (**A**, **B**) Subcellular localization of RipV1 (A) and RLCK-IXb-1 (B). RipV1-YFP or RLCK-IXb-1-YFP were co-expressed with AtFLS2- mCherry in *N. benthamiana*. AtFLS2-mCherry was used as a plasma membrane marker. Subcellular localization was observed by confocal microscopy. Fluorescence images were merged with brightfield (BF) images. Scale bars represent 50 µm. Fluorescence intensity of YFP and mCherry across the section indicated by the white dotted line is shown on the right. (**C**) Protein expression of YFP-tagged RipV1 and RLCK-IXb-1 in *N. benthamiana*. RipV1-YFP and RLCK-IXb-1-YFP were transiently expressed in *N. benthamiana* by agroinfiltration. Leaf samples were harvested 45 hours post infiltration. Total protein extracts were probed with anti-GFP antibody. Ponceau S (PS) staining was used to verify equal loading and protein transfer. The red asterisk marks the position of the RLCK-IXb-1-YFP fusion protein.

**Figure S4.** Alignment of kinase domains in AtPUB32, AtPUB33, and their homologs from tomato and *N. benthamiana*. Conserved regions corresponding to key kinase catalytic motifs including the glycine-rich loop (GxGxxG), the ATP-binding motif (VAIK), the catalytic loop (HRD) and the activation segment (DFG) according to Roux et al. (2014) are framed by red boxes. Underlined amino acids denote key residues for function. Alignment was performed using ClustalW, and the image was generated using ESPript 3.0 with similarity-based coloring. Fully conserved residues (100% identity) are shown with white bold text on a black background. Positions with ≥70% similarity and a global conservation score ≥0.7 are shown in bold black without shading. Residue numbering corresponds to the full-length AtPUB33 protein (At2G45910) used as the reference sequence for alignment.

**Figure S5.** RLCK-IXb-1 cannot trans-phosphorylate RipV1 *in vitro*. Recombinant RLCK-IXb-1 derivatives (RLCK-IXb-1, RLCK-IXb-1^D659A^, RLCK-IXb-1KDm, and RLCK-IXb-1KDm^D659A^) fused with N-terminal MBP tag and C-terminal FLAG tag were incubated with RipV1. SlPti1b was used as an auto-phosphorylation control. Phosphorylation reaction using γ^32^P-ATP was detected by autoradiography (Autorad) following SDS-PAGE. Coomassie Brilliant Blue (CBB) staining was performed to verify protein loading. Black asterisks indicate the positions of the recombinant RLCK-IXb-1 derivatives and SlPti1b. Blue asterisks indicate the position of the recombinant RipV1 fused with C-terminal Myc tag.

**Figure S6.** RipV1 has an E3 ubiquitin ligase activity *in vitro*. RipV1 and RipV1^C452A^ recombinant proteins were incubated with HA-Ubiquitin in the presence or absence of human E1 (UBE1) and E2 (UbcH5B). Proteins in the mixtures were probed with anti-HA and anti-Myc antibodies following separation by SDS-PAGE.

**Figure S7.** Relative expression of *NbRLCK-IXb-1*, *-2*, *-3*, and *-9* in silenced *N. benthamiana* plants. Silencing efficiency for *NbRLCK-IXb-1* **(A)**, *-2* **(B)**, *-3* **(C)**, and *-9* **(D)** in TRV:*NbRLCK-IXb- 1/2/3/9* plants. The silenced genes and their corresponding accession numbers are as follows: *NbRLCK-IXb-1-1* (*Niben261Chr03g0421011.1*), *NbRLCK-IXb-1-2* (*Niben261Chr09g1137026.1*), *NbRLCK-IXb-2-1* (*Niben261Chr06g0821015.1*), *NbRLCK-IXb-2-2* (*Niben261Chr08g0471021.1*), *NbRLCK-IXb-3-1* (*Niben261Chr06g0936005.1*), *NbRLCK-IXb-3-2* (*Niben261Chr08g0547001.1*), *NbRLCK-IXb-9-1* (*Niben261Chr12g0682001.1*), and *NbRLCK-IXb-9-2* (*Niben261Chr19g0053013.1*). Silencing was evaluated by quantifying gene expression in *N. benthamiana* using qRT-PCR with primers listed in Table S2. Each graph shows gene expression normalized by *NbEF1α* and relative to the expression in TRV:EV plants. Data are presented as mean ± standard deviation. Asterisks indicate statistically significant differences compared to TRV:EV plants. The differences were analyzed by unpaired *t*-test (∗∗∗∗, *P* < 0.0001; ∗∗∗, *P* < 0.001; ∗∗, *P* < 0.01 and ns, non-significant).

**Figure S8.** RipV1-induced cell death occurs in *N. benthamiana* plants impaired for effector recognition. (**A**) RipV1-induced cell death was delayed in *NbSGT1* silenced plants. FLAG-GFP, RipV1-FLAG, RipE1-FLAG, and RipA1-FLAG were transiently expressed in TRV:EV or TRV:*NbSGT1* plants. Leaves were photographed 3 days post infiltration (dpi). False-color images representing quantum yield (QY) were generated using FluorCam software. A customized color scale was applied to enhance visual contrast, in which black regions indicate areas of cell death. (**B**) Box-and-whisker plot shows QY values from three independent repeats (n = 25) of the experiment shown in (A). Boxes indicate interquartile range, median is shown by the central line, whiskers by the minimum and maximum of QY, and ‘+’ denotes the mean. Different letters indicate statistically significant differences analyzed by two-way analysis of variance (ANOVA) followed by Tukey’s multiple comparisons test (*P* < 0.0001). (**C**) Protein accumulation of FLAG-GFP or RipV1-FLAG in TRV:EV and TRV:*NbSGT1* plants. Total protein extracts were probed with anti-FLAG-HRP antibodies. Ponceau S (PS) staining was used to verify equal loading and protein transfer. (**D**) RipV1-induced cell death is independent of NLR network signaling pathways. GFP, RipV1^C452A^-FLAG, RipV1-FLAG, XopQ-FLAG, AVRblb2, and Rpi-blb2 were expressed in wild-type, *epss* (*EDS1*, *PAD4*, *SAG101a*, and *SAG101b*), *nrg1* and *nrc2/3/4 N. benthamiana* knock-out lines. Leaves were photographed 3dpi. False-color images representing QY were generated using FluorCam software. A customized color scale was applied to enhance visual contrast, in which black regions indicate areas of cell death. (**E**) Box-and-whisker plot shows QY values from 3 independent repeats (n = 30) of the experiment shown in (D). Boxes indicate interquartile range, median is shown by the central line, whiskers by the minimum and maximum of QY, and ‘+’ denotes the mean. Different letters indicate statistically significant differences analyzed by two-way ANOVA followed by Tukey’s multiple comparisons test (*P* < 0.0001).

**Figure S9.** RipV1 and RLCK-IXb-1 protein accumulation. **(A)** Accumulation of RipV1 after agroinfiltration with different OD_600_ values in TRV:EV and TRV:*NbRLCK-IXb-1/2/3/9* plants. Leaf samples were harvested at 35 hours post-infiltration (hpi). Proteins from total protein extracts were probed with anti-FLAG-HRP antibodies. Ponceau S (PS) staining was used to verify equal loading and protein transfer. **(B)** Accumulation of GFP, RipV1-FLAG, RLCK-IXb-1-HA, and RLCK-IXb-2-HA after agroinfiltration in *N. benthamiana*. Leaf samples were harvested at 35 hpi. Total protein extracts were probed with anti-FLAG-HRP, anti-HA, anti-GFP, or anti-Actin antibodies.

**Table S1.** Prediction of RLCK-IXb-1 palmitoylation sites. Table S2. Primers used in this study.

