## Supplementary material for "The *Ralstonia solanacearum* E3 ligase effector RipV1 targets subfamily IXb receptor-like cytoplasmic kinases that negatively regulate immunity in *Nicotiana benthamiana*": Choi J et al_supp info

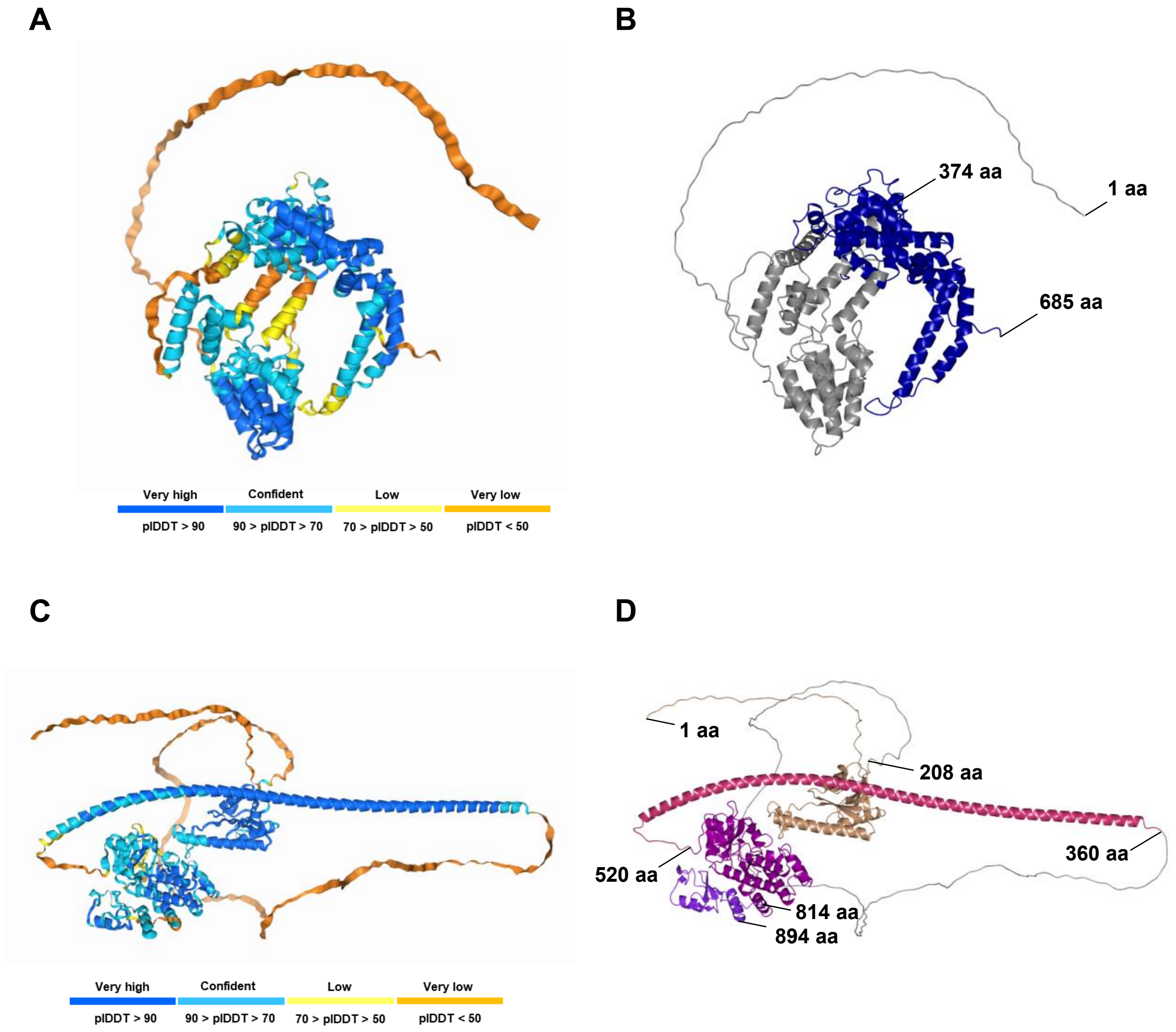


**Figure S1. 3D structure models of RipV1 and RLCK-IXb-1. (A, B)** RipV1 protein structure with predicted local distance difference test (plDDT) confidence score (A) and colored domains (B). RipV1 truncation constructs used in yeast two-hybrid assays are shown in (B). The grey region (1-374 aa) corresponds to the N-terminal domain, and the navy region (374-685 aa) includes the NEL domain (374-604 aa; conserved domain database (CDD) cl38455). **(C, D)** RLCK-IXb-1 protein structure with plDDT confidence score (C) and colored regions (D). RLCK-IXb-1 domains were defined according to the domain analysis of RLCK-IXb-1: USP (1-208 aa; brown; cd01989), Smc (361-520 aa; maroon; cl34310), Kinase (521-814 aa; dark magenta; cl21453), and PUB (815-894 aa; violet; cd16655) domains. The unfolded region (209-360 aa) between USP and Smc domains is colored in grey. Domain analysis was conducted by NCBI CD-search and 3D structure was predicted by AlphaFold 3.

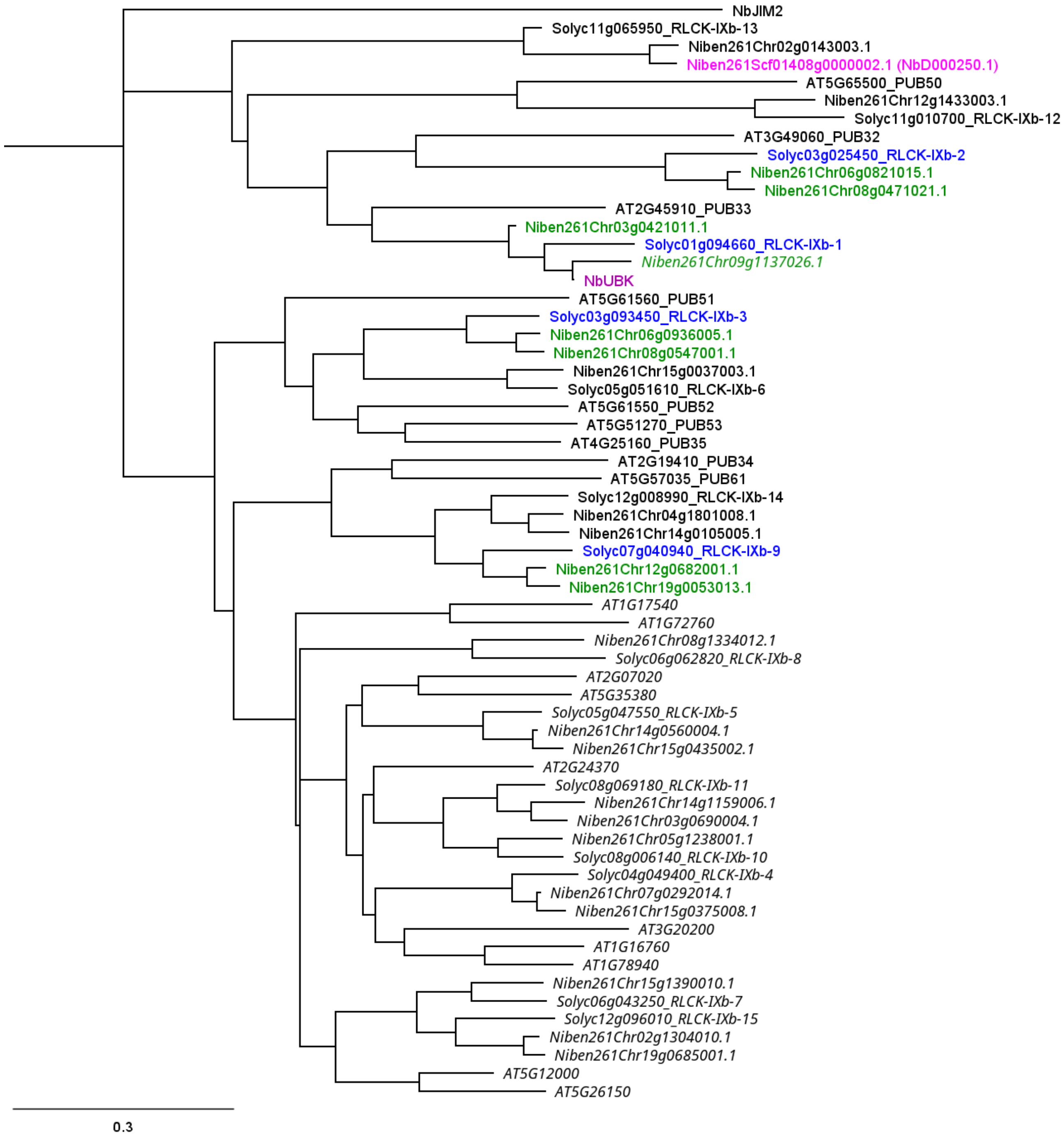


**Figure S2. Phylogenetic analysis of RLCK subfamily IXb in Arabidopsis, tomato, and *N. benthamiana*.** Arabidopsis and tomato RLCK-IXb protein sequences were obtained from Sakamoto et al. (2012). RLCK-IXb members were retrieved from the *N. benthamiana* genome (v2.6.1) available at (https://solgenomics.net/organism/Nicotiana_benthamiana/genome) using the 18 Arabidopsis and 15 tomato RLCK-IXb protein sequences as queries. The RLCK-XII XOPJ4 IMMUNITY 2 (JIM2) was used as an outgroup (Schultink et al., 2019). The tree was built on full-length amino acid sequences using the neighbor-joining method with 1,000 bootstrap replicates performed by the Jukes-Cantor model. The scale bar indicates evolutionary distance. RLCKs interacting with RipV1 in yeast two-hybrid are highlighted in blue. *N. benthamiana* homologs of RLCK-IXb-1, -2, -3, and -9 targeted by virus-induced gene silencing are highlighted in green. The protein highlighted in magenta is identical to NbD000250.1 reported as AtPUB33 homolog by Dong et al. (2023). Another AtPUB33 homolog, NbUBK, described in He et al. (2018) is labeled in purple. Members of RLCK subfamily IXb that lack a PUB domain are shown in italics.

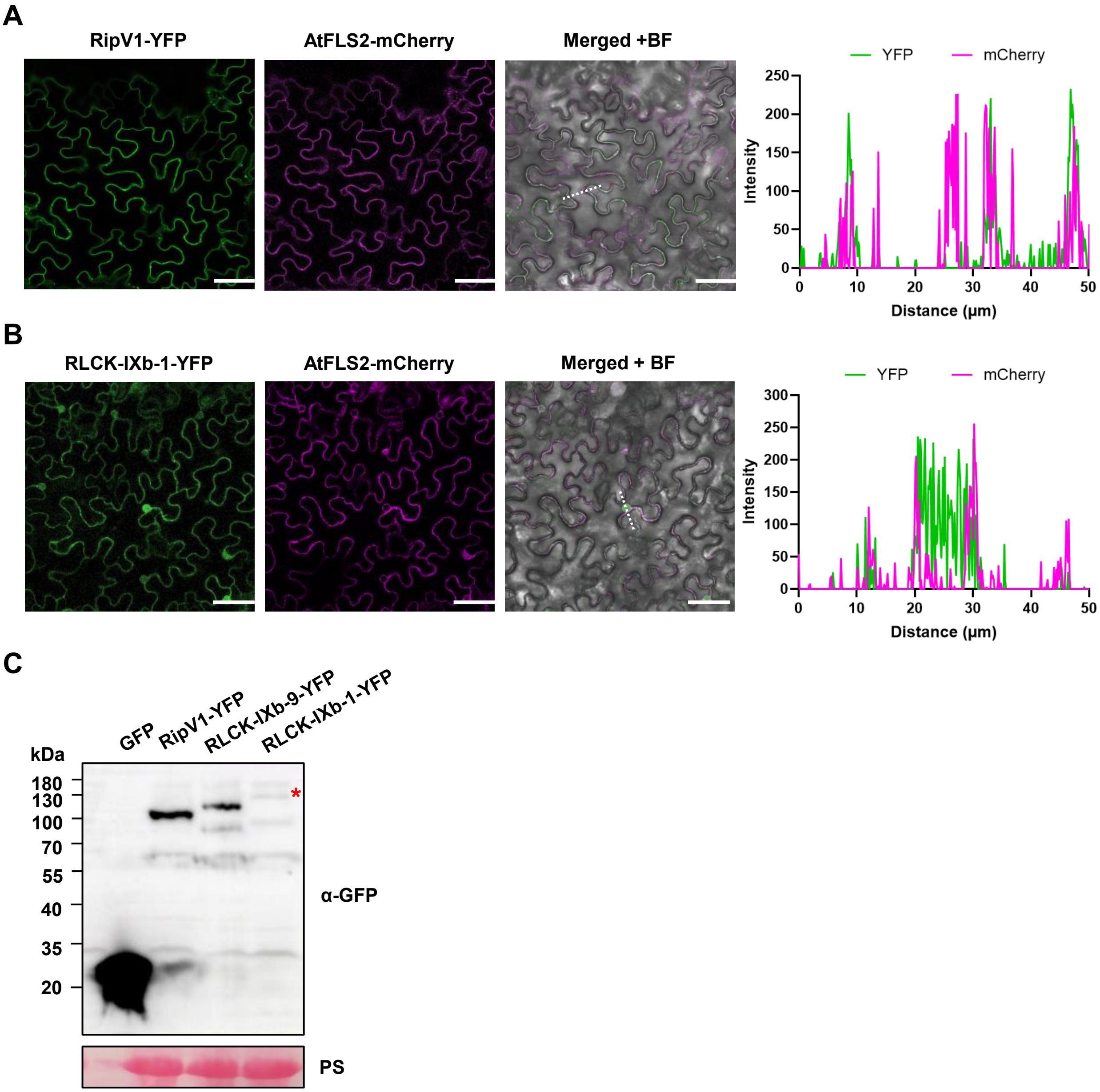


**Figure S3. Subcellular localization of RipV1 and RLCK-IXb-1.** (**A**, **B**) Subcellular localization of RipV1 (A) and RLCK-IXb-1 (B). RipV1-YFP or RLCK-IXb-1-YFP were co-expressed with AtFLS2-mCherry in *N. benthamiana*. AtFLS2-mCherry was used as a plasma membrane marker. Subcellular localization was observed by confocal microscopy. Fluorescence images were merged with brightfield (BF) images. Scale bars represent 50 µm. Fluorescence intensity of YFP and mCherry across the section indicated by the white dotted line is shown on the right. (**C**) Protein expression of YFP-tagged RipV1 and RLCK-IXb-1 in *N. benthamiana*. RipV1-YFP and RLCK-IXb-1-YFP were transiently expressed in *N. benthamiana* by agroinfiltration. Leaf samples were harvested 45 hours post infiltration. Total protein extracts were probed with anti-GFP antibody. Ponceau S (PS) staining was used to verify equal loading and protein transfer. The red asterisk marks the position of the RLCK-IXb-1-YFP fusion protein.

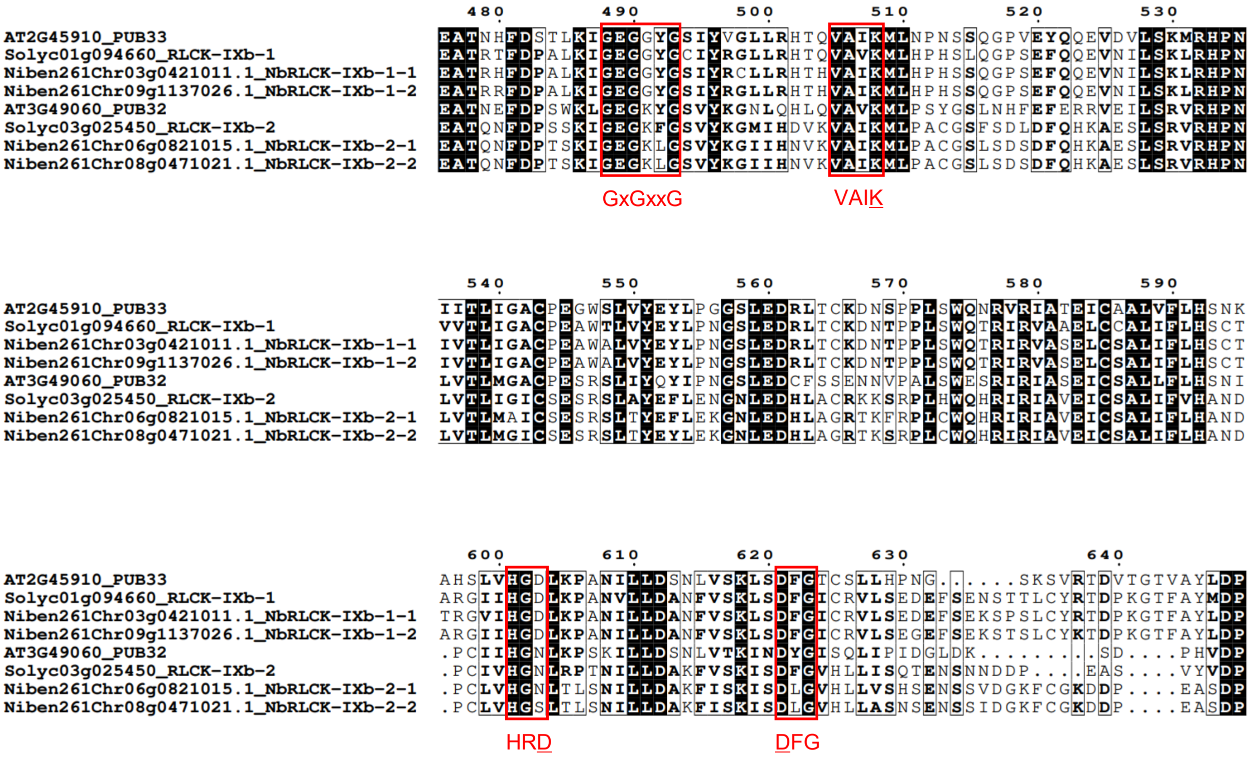


**Figure S4. Alignment of kinase domains in AtPUB32, AtPUB33, and their homologs from tomato and *N. benthamiana*.** Conserved regions corresponding to key kinase catalytic motifs including the glycine-rich loop (GxGxxG), the ATP-binding motif (VAIK), the catalytic loop (HRD) and the activation segment (DFG) according to Roux et al. (2014) are framed by red boxes. Underlined amino acids denote key residues for function. Alignment was performed using ClustalW, and the image was generated using ESPript 3.0 with similarity-based coloring. Fully conserved residues (100% identity) are shown with white bold text on a black background. Positions with ≥70% similarity and a global conservation score ≥0.7 are shown in bold black without shading. Residue numbering corresponds to the full-length AtPUB33 protein (At2G45910) used as the reference sequence for alignment.

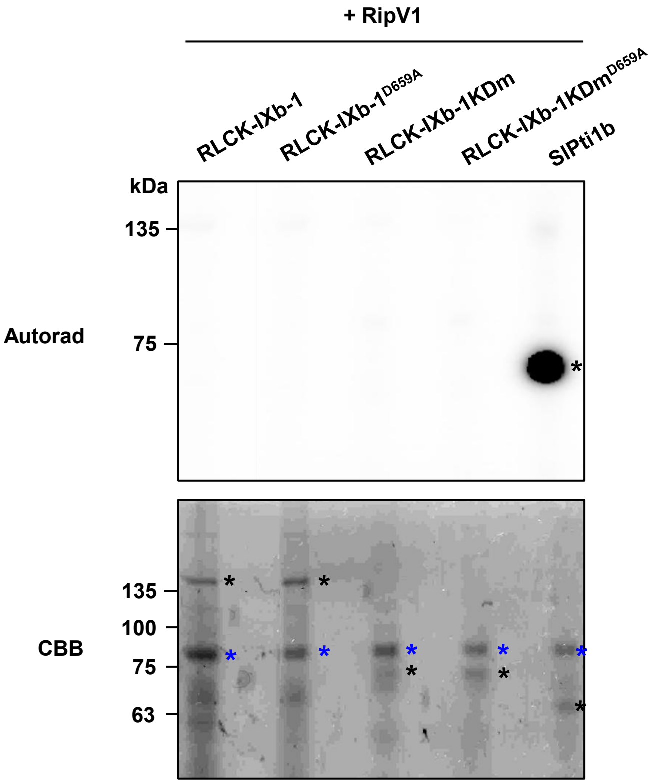


**Figure S5. RLCK-IXb-1 cannot trans-phosphorylate RipV1 *in vitro*.** Recombinant RLCK-IXb-1 derivatives (RLCK-IXb-1, RLCK-IXb-1$^{D659A}$, RLCK-IXb-1KDm, and RLCK-IXb-1KDm$^{D659A}$) fused with N-terminal MBP tag and C-terminal FLAG tag were incubated with RipV1. SlPti1b was used as an auto-phosphorylation control. Phosphorylation reaction using g$^{32}$P-ATP was detected by autoradiography (Autorad) following SDS-PAGE. Coomassie Brilliant Blue (CBB) staining was performed to verify protein loading. Black asterisks indicate the positions of the recombinant RLCK-IXb-1 derivatives and SlPti1b. Blue asterisks indicate the position of the recombinant RipV1 fused with C-terminal Myc tag.

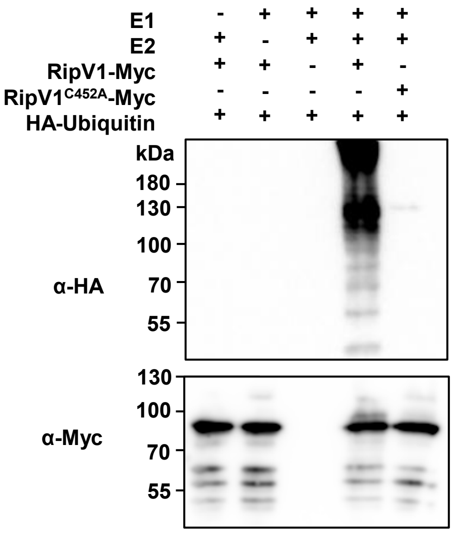


**Figure S6. RipV1 has an E3 ubiquitin ligase activity *in vitro*.** RipV1 and RipV1$^{C452A}$ recombinant proteins were incubated with HA-Ubiquitin in the presence or absence of human E1 (UBE1) and E2 (UbcH5B). Proteins in the mixtures were probed with anti-HA and anti-Myc antibodies following separation by SDS-PAGE.

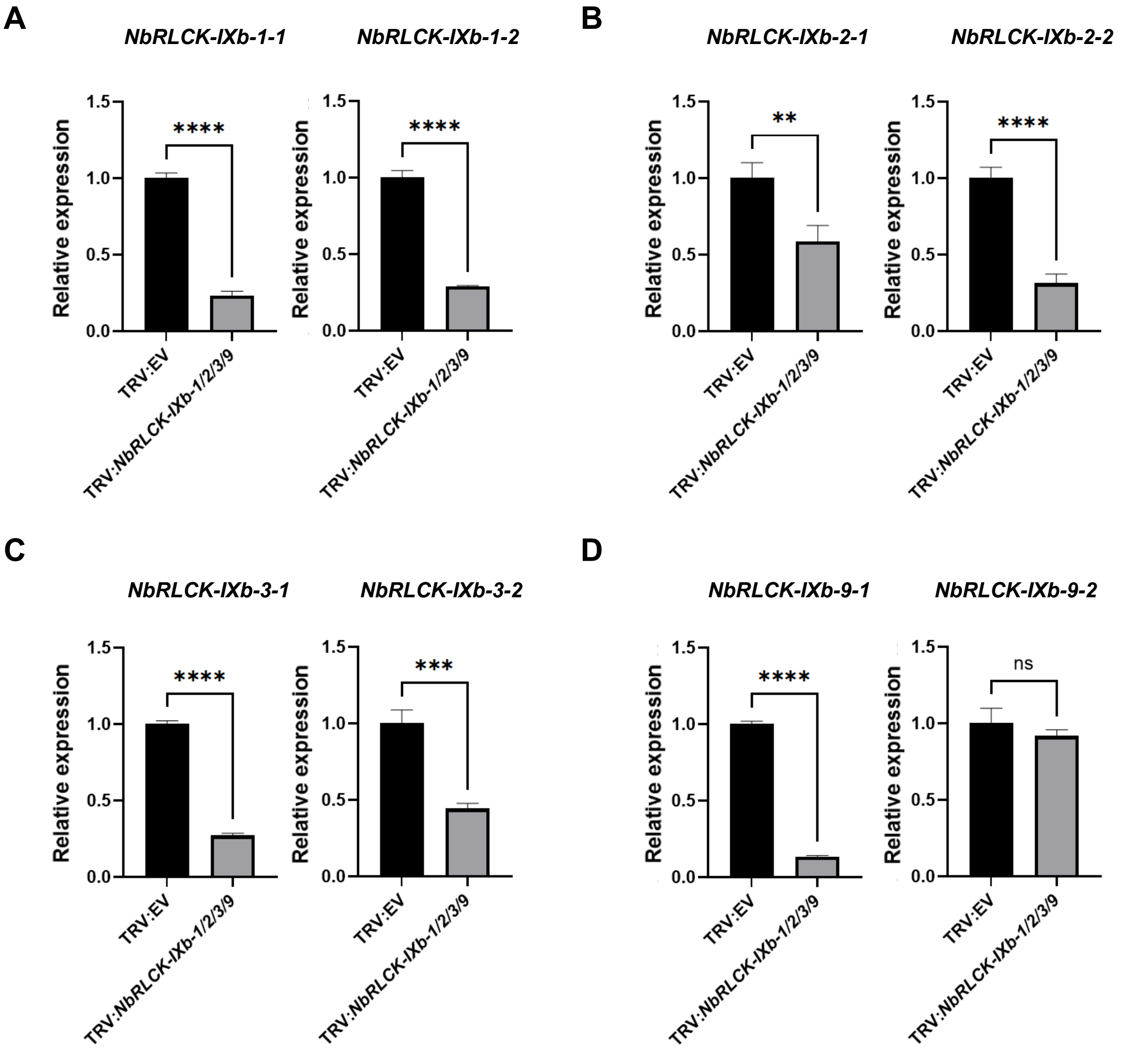


**Figure S7. Relative expression of *NbRLCK-IXb-1*, *-2*, *-3*, and *-9* in silenced *N. benthamiana* plants.** Silencing efficiency for *NbRLCK-IXb-1* **(A)**, *-2* **(B)**, *-3* **(C)**, and *-9* **(D)** in TRV:*NbRLCK-IXb-1/2/3/9* plants. The silenced genes and their corresponding accession numbers are as follows: *NbRLCK-IXb-1-1* (*Niben261Chr03g0421011.1*), *NbRLCK-IXb-1-2* (*Niben261Chr09g1137026.1*), *NbRLCK-IXb-2-1* (*Niben261Chr06g0821015.1*), *NbRLCK-IXb-2-2* (*Niben261Chr08g0471021.1*), *NbRLCK-IXb-3-1* (*Niben261Chr06g0936005.1*), *NbRLCK-IXb-3-2* (*Niben261Chr08g0547001.1*), *NbRLCK-IXb-9-1* (*Niben261Chr12g0682001.1*), and *NbRLCK-IXb-9-2* (*Niben261Chr19g0053013.1*). Silencing was evaluated by quantifying gene expression in *N. benthamiana* using qRT-PCR with primers listed in Table S2. Each graph shows gene expression normalized by *NbEF1α* and relative to the expression in TRV:EV plants. Data are presented as mean ± standard deviation. Asterisks indicate statistically significant differences compared to TRV:EV plants. The differences were analyzed by unpaired *t*-test (****, $P < 0.0001$; ***, $P < 0.001$; **, $P < 0.01$ and ns, non-significant).

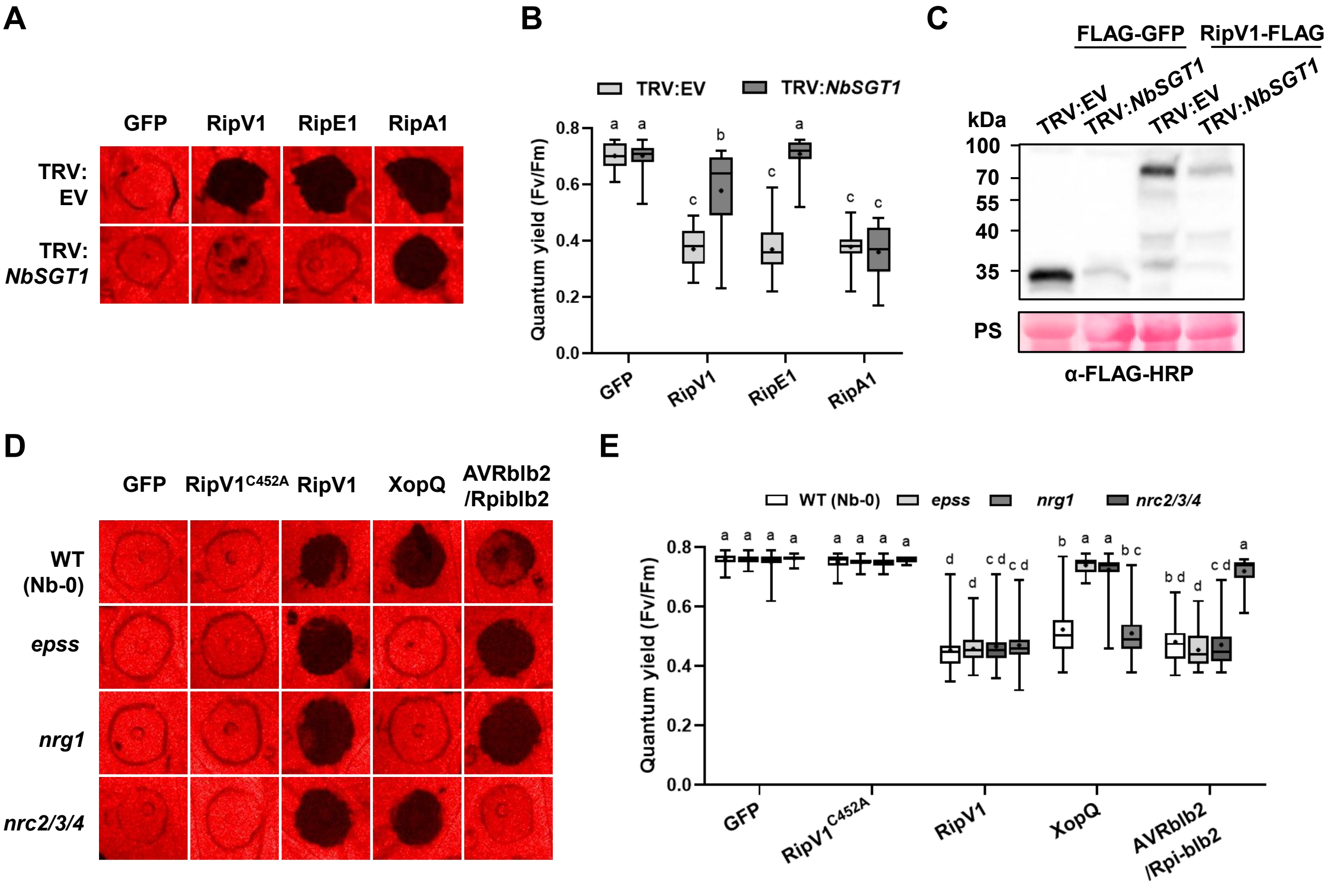


**Figure S8. RipV1-induced cell death occurs in *N. benthamiana* plants impaired for effector recognition.** (**A**) RipV1-induced cell death was delayed in *NbSGT1* silenced plants. FLAG-GFP, RipV1-FLAG, RipE1-FLAG, and RipA1-FLAG were transiently expressed in TRV:EV or TRV:*NbSGT1* plants. Leaves were photographed 3 days post infiltration (dpi). False-color images representing quantum yield (QY) were generated using FluorCam software. A customized color scale was applied to enhance visual contrast, in which black regions indicate areas of cell death. (**B**) Box-and-whisker plot shows QY values from three independent repeats (n = 25) of the experiment shown in (A). Boxes indicate interquartile range, median is shown by the central line, whiskers by the minimum and maximum of QY, and '+' denotes the mean. Different letters indicate statistically significant differences analyzed by two-way analysis of variance (ANOVA) followed by Tukey's multiple comparisons test ($P < 0.0001$). (**C**) Protein accumulation of FLAG-GFP or RipV1-FLAG in TRV:EV and TRV:*NbSGT1* plants. Total protein extracts were probed with anti-FLAG-HRP antibodies. Ponceau S (PS) staining was used to verify equal loading and protein transfer. (**D**) RipV1-induced cell death is independent of NLR network signaling pathways. GFP, RipV1$^{C452A}$-FLAG, RipV1-FLAG, XopQ-FLAG, AVRblb2, and Rpi-blb2 were expressed in wild-type, *epss* (*EDS1*, *PAD4*, *SAG101a*, and *SAG101b*), *nrg1* and *nrc2/3/4* *N. benthamiana* knock-out lines. Leaves were photographed 3dpi. False-color images representing QY were generated using FluorCam software. A customized color scale was applied to enhance visual contrast, in which black regions indicate areas of cell death. (**E**) Box-and-whisker plot shows QY values from 3 independent repeats (n = 30) of the experiment shown in (D). Boxes indicate interquartile range, median is shown by the central line, whiskers by the minimum and maximum of QY, and '+' denotes the mean. Different letters indicate statistically significant differences analyzed by two-way ANOVA followed by Tukey's multiple comparisons test ($P < 0.0001$).

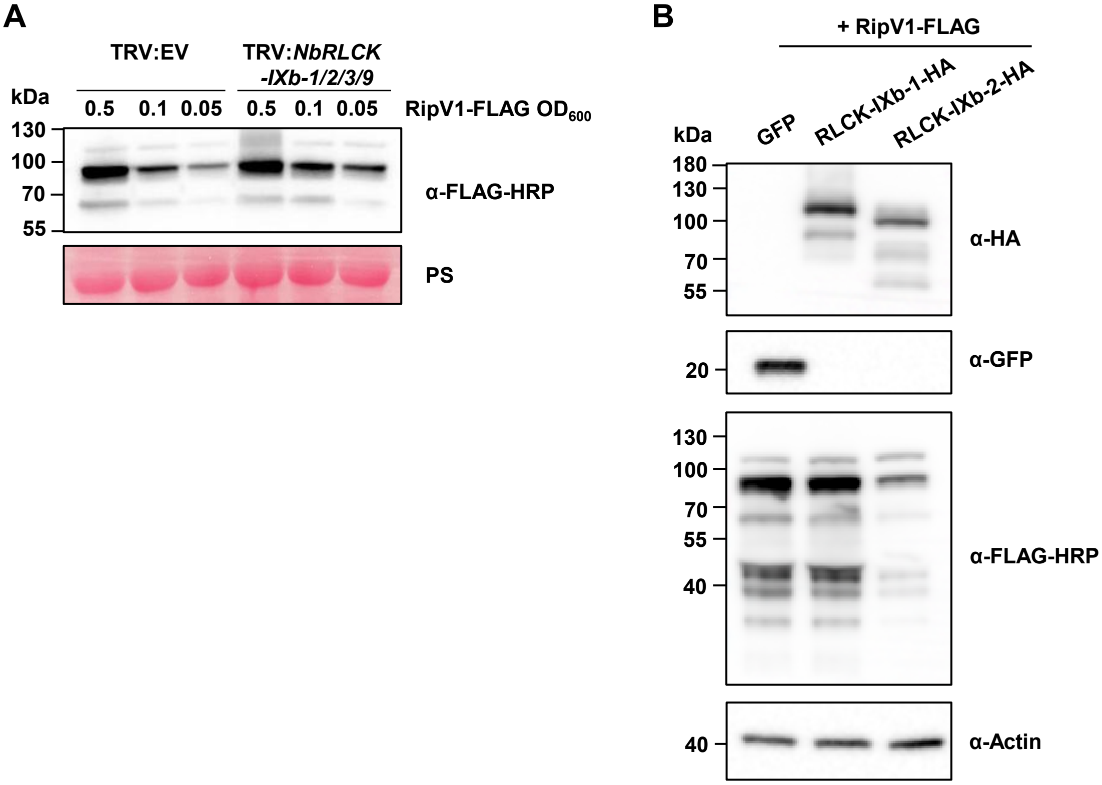


**Figure S9. RipV1 and RLCK-IXb-1 protein accumulation. (A)** Accumulation of RipV1 after agroinfiltration with different $OD_{600}$ values in TRV:EV and TRV:*NbRLCK-IXb-1/2/3/9* plants. Leaf samples were harvested at 35 h post-infiltration. Proteins from total protein extracts were probed with anti-FLAG-HRP antibodies. Ponceau S (PS) staining was used to verify equal loading and protein transfer. **(B)** Accumulation of GFP, RipV1-FLAG, RLCK-IXb-1-HA, and RLCK-IXb-2-HA after agroinfiltration in *N. benthamiana*. Leaf samples were harvested at 35 h post-infiltration. Total protein extracts were probed with anti-FLAG-HRP, anti-HA, anti-GFP, or anti-Actin antibodies.

**Table S1. Prediction of RLCK-IXb-1 palmitoylation sites.**

| Position (aa) | Peptide sequence | Prediction resource |
|---|---|---|
| 75 | ALHKSGGRKI-**C**-IVHVHTPAQK | $PTMGPT_2$[a] |
| 125 | HMILEKYILI-**C**-GRAGVCADKL | $PTMGPT_2$; GPS-Palm[b] |
| 167 | KLVMGAAANK-**C**-YSKKMSDLRS | GPS-Palm |
| 192 | YVRLQAPTFC-**C**-ICFVCKGNLI | $PTMGPT_2$; GPS-Palm |
| 197 | APTFCCICFV-**C**-KGNLIFTRES | $PTMGPT_2$ |
| 550 | ALKIGEGGYG-**C**-IYRGLLRHTQ | GPS-Palm |
| 783 | AKQLAHLAMS-**C**-CDKNSRCRPE | GPS-Palm |
| 784 | KQLAHLAMSC-**C**-DKNSRCRPEL | GPS-Palm |
| 809 | WKVLEPMRAS-**C**-GASSFRIDSE | GPS-Palm |

[a] https://nsclbio.jbnu.ac.kr/tools/ptmgpt2
[b] https://gpspalm.biocuckoo.cn; prediction was performed using the 'high' threshold setting

### Table S2. Primers used in this study.

| Primer | Sequence | Description |
|---|---|---|
| RipV1-C452A-F | GACTGCACTCGGTACTGCCGGAGATAATGTTGCAGA | Site-directed mutagenesis |
| RipV1-C452A-R | TCTGCAACATTATCTCCGGCAGTACCGAGTGCAGTC | Site-directed mutagenesis |
| RipV1-NT-F | GGTCTCA AATGCCGACCCGTGTTCCA | Golden Gate Cloning |
| RipV1-NT-R | GGTCTCA CGAATTGCTTGATCTCTGCAGCCA | Golden Gate Cloning |
| RipV1-NEL-F | GGTCTCA AATGTGGTTAAAGCTAGCGGATATGA | Golden Gate Cloning |
| RipV1-NEL-R | GGTCTCA CGAAACGGCTCCCCTGGCTTGG | Golden Gate Cloning |
| RLCK-IXb-1-D659A-F | GGGGCATAATTCATGGTGCTCTAAAACCAGCAAATG | Site-directed mutagenesis |
| RLCK-IXb-1-D659A-R | CATTTGCTGGTTTTAGAGCACCATGAATTATGCCCC | Site-directed mutagenesis |
| RLCK-IXb-1-C830A-F | CCCATCCTATTTTATTGCTCCCATATTTCAGG | Site-directed mutagenesis |
| RLCK-IXb-1-C830A-F | CCTGAAATATGGGAGCAATAAAATAGGATGGG | Site-directed mutagenesis |
| RLCK-IXb-1_1-208_F | GGTCTCAAATGGCTTTGGAGACTCCATCATC | Golden Gate Cloning |
| RLCK-IXb-1_1-208_R | GGTCTCACGAACTTACTTTCTCTTGTAAAAATGAGATTCCCTTTG | Golden Gate Cloning |
| RLCK-IXb-1_209-360-F | GGTCTCAAATGTCAGACAGACTTAATACAGACAGTGTG | Golden Gate Cloning |
| RLCK-IXb-1_209-360-R | GGTCTCACGAAGCTTCCAGCTAGCTCGTTATTC | Golden Gate Cloning |
| RLCK-IXb-1_361-520-F | GGTCTCAAATGAACGATGAACTCTATGATAGATATGAGCA | Golden Gate Cloning |
| RLCK-IXb-1_361-520-R | GGTCTCACGAATGAAGATGTAGATGTTAATGAAGACCC | Golden Gate Cloning |
| RLCK-IXb-1_521-814-F | GGTCTCAAATGCTATTTGCTGAATTTTATTTTCATGAAATTGAAGAAGC | Golden Gate Cloning |
| RLCK-IXb-1_521-814-R | GGTCTCACGAAAAAAGATGATGCCCCACATGAAGC | Golden Gate Cloning |
| RLCK-IXb-1_815-894-F | GGTCTCAAATGAGAATAGATTCCGAAGAGCATTGTG | Golden Gate Cloning |
| RLCK-IXb-1_815-894-R | GGTCTCACGAAATTCTGTTGCAGCCACTCCTGAA | Golden Gate Cloning |
| NbRLCK-IXb-1-1_VIGS-F | GGTCTCAATGACTGGAGAGCCAACTCTTAAATTCTGAC | VIGS |
| NbRLCK-IXb-1-1_VIGS-R | GGTCTCAATAGCAGCGCCTCAGCTATATTGAGTG | VIGS |
| NbRLCK-IXb_2-VIGS-F | GGTCTCACTATGATGAAGATTCAGCATGACCTGTGC | VIGS |
| NbRLCK-IXb_2-VIGS-R | GGTCTCACTTCAAAGATATCAGAAGCTCCACTGCTTG | VIGS |
| NbRLCK-IXb_3-VIGS-F | GGTCTCAGAAGTCCAGTTTACTAACAGAAAAGTCAGACCTG | VIGS |
| NbRLCK-IXb_3-VIGS-R | GGTCTCAAAGCGACGGTTTCATTTGAGATTGCTCTTCT | VIGS |
| NbRLCK-IXb_9-VIGS-F | GGTCTCAGGAGAAGAATGCGCTTAGAATTACAAAATACCATTG | VIGS |
| NbRLCK-IXb_9-VIGS-R | GGTCTCATCATCTTTCTTCTCTTTTCTGGGCTGCA | VIGS |
| NbRLCK-IXb-1-1_qPCR-F | GGCAGAGGAGAGACATAGAGG | qPCR |
| NbRLCK-IXb-1-1_qPCR-R | GTTGTGTCAGAATTTAAGAGTTGGCT | qPCR |
| NbRLCK-IXb-1-2_qPCR-F | GTGGCCGACGATATGATATATGTTG | qPCR |
| NbRLCK-IXb-1-2_qPCR-R | TGCACATCCAACTGATCTATGTTAAAC | qPCR |
| NbRLCK-IXb-2-1_qPCR-F | GTTTGTAGCAGTGGGAAAGAATGTG | qPCR |
| NbRLCK-IXb-2-1_qPCR-R | CAGCTTTGTACCTGAAACCTTCC | qPCR |
| NbRLCK-IXb-2-2_qPCR-F | TTACTCAAAACAATTGTCAGAGCTCAAG | qPCR |
| NbRLCK-IXb-2-2_qPCR-R | GACAATGACTCTTTGCTAGTTAAATGAGA | qPCR |
| NbRLCK-IXb-3-1_qPCR-F | GAATCCATTAAACTCTTCCACCCTTG | qPCR |
| NbRLCK-IXb-3-1_qPCR-R | CATCCCGCACTTGTGATATAGGT | qPCR |
| NbRLCK-IXb-3-2_qPCR-F | ATGTGAATCCATTAAACTCTTCCACC | qPCR |
| NbRLCK-IXb-3-2_qPCR-R | CTTGTGAAATAGGTATAGAGTTCCCCA | qPCR |
| NbRLCK-IXb-9-1_qPCR-F | GTGGGCTATAGAGAAGTTGTTGC | qPCR |
| NbRLCK-IXb-9-1_qPCR-R | GCACCACAGTTTCAACACTTCTTC | qPCR |
| NbRLCK-IXb-9-2_qPCR-F | GATGACAATGTGGTGGAAATGTATATTG | qPCR |
| NbRLCK-IXb-9-2_qPCR-R | CAACACTAGAGTATTAATCCCAGCG | qPCR |
| NbEF1α-qPCR-F | TGGACACAGGGACTTCATCA | qPCR |
| NbEF1α-qPCR-R | CAAGGGTGAAAGCAAGCAAT | qPCR |
| pLexA-vector-F | ACCAATTGTCGTAGATCTT | PCR to confirm presence of plasmid |
| pLexA-vector-R | CATAAATCATAAGAAATTCG | PCR to confirm presence of plasmid |
| pB42-AD-vector-F | GGATGTTAACGATACCAG | PCR to confirm presence of plasmid |
| pB42-AD-vector-R | CATCTTTTCGTAAATTTCT | PCR to confirm presence of plasmid |
